# Proton receptors regulate synapse-specific reconsolidation in the amygdala

**DOI:** 10.1101/2021.01.04.425235

**Authors:** Erin E Koffman, Charles M Kruse, Kritika Singh, FarzanehSadat Naghavi, Jennifer Egbo, Sandra Boateng, Mark Houdi BA, Boren Lin, Jacek Debiec, Jianyang Du

## Abstract

When an extinction procedure is performed within the reconsolidation window, the original aversive memory can be replaced by one that is less traumatic. Recent studies revealed that carbon dioxide (CO_2_) inhalation during retrieval enhances memory lability. However, the effects of CO_2_ inhalation on the central nervous system can be extensive, and there is lack of evidence suggesting that the effects of CO_2_ are selective to a reactivated memory. We discovered that CO_2_ inhalation paired with memory retrieval potentiates the specific aversive memory trace, resulting in greater memory lability. The specific effects of CO_2_ depend on acid-sensing ion channels (ASICs), the proton receptors that are involved in synaptic transmission and plasticity in the amygdala. In addition, CO_2_ inhalation alters memory lability via synaptic plasticity at selectively targeted synapses. Overall, our results suggest that inhaling CO_2_ during the retrieval event increases the lability of an aversive memory through a synapse-specific reconsolidation process.

## INTRODUCTION

Recently, aversive memory research in both rodents and humans has focused on a window of time after aversive memory retrieval, in which the memory is labile and subject to intervention (Kida, 2020; Lee, 2009; Monfils et al., 2009; Nader et al., 2000; Sara, 2000; Schiller et al., 2010; Tronson and Taylor, 2007). This window of time is known as a reconsolidation window, and is thought to last up to six hours after initiation (Clem and Huganir, 2010; Duvarci and Nader, 2004; Monfils et al., 2009; Schiller et al., 2010). Several studies have demonstrated that interrupting the updating process aroused by retrieval prevents memory restorage, generating selective amnesia (Agren et al., 2012; Clem and Huganir, 2010; Mactutus et al., 1979; Monfils et al., 2009; Nader et al., 2000; Schiller et al., 2010). Studies using rodent models have indicated that pharmacological intervention within the reconsolidation window successfully erases reactivated specific aversive memory (Nader et al., 2000; Sara and Hars, 2006; Schiller et al., 2010). Despite the efficacy, ethical and practical concerns may prevent similar pharmacological interventions from being used in humans (Agren et al., 2012; Schiller et al., 2010). Recently, drug-free paradigms have been proposed that effectively prevent the return of aversive memories in both rodents and humans (Bjorkstrand et al., 2016; Clem and Huganir, 2010; Huang et al., 2020; Liu et al., 2014; Monfils et al., 2009; Schiller et al., 2010). Monfils *et al*. developed a novel protocol in which an isolated retrieval trial was followed by an extinction event within a specific time frame. This resulted in the weakening of the original memory trace, thereby preventing reversion of the original aversive memory by spontaneous recovery, renewal, or reinstatement (Clem and Huganir, 2010; Jarome et al., 2015; Monfils et al., 2009; Quirk et al., 2010; Wu et al., 2017). These studies suggest the mechanism by which retrieval changes the lability of memory may become a target for novel clinical treatments for anxiety disorders.

Currently, the outcomes of the combination of retrieval and extinction paradigms are variable, depends on multiple factors, in both rodents (Auber et al., 2013; Chan et al., 2010; Costanzi et al., 2011; Goode et al., 2017; Ishii et al., 2012) and humans (Golkar et al., 2012; Klucken et al., 2016). Because it is difficult to completely erase the original memory, a more reliable strategy for triggering memory erasure is necessary. In search of this strategy, we studied the effects that CO_2_ inhalation may have on the erasure of a specific memory. As we showed previously, when mice receive a retrieval tone supplemented by CO_2_ inhalation, the memory becomes more labile than the presentation of a retrieval tone alone (Du et al., 2017). Within the reconsolidation window, the labile memory becomes more convertible, either weakened by extinction or strengthened by reconditioning. Moreover, the effects of CO_2_ inhalation on memory lability were dependent on acid-sensing ion channels (ASICs), since disrupting ASICs in the amygdala eliminated these effects of CO_2_ (Du et al., 2017).

Recent studies have revealed that protons are potential neurotransmitters (Du et al., 2014; Highstein et al., 2014; Kawasaki et al., 2009) and ASICs serve as postsynaptic proton receptors that play key roles in neurotransmission and synaptic plasticity in the amygdala, a brain region that is critical for the formation of aversive memories (Cohen et al., 2017; Du et al., 2014; Farley et al., 2018; Ziemann et al., 2009). ASICs are members of the Degenerin/ Epithelial sodium channel (DEG/ ENaC) family (Ben-Shahar, 2011). To date, six family members have been identified (ASIC1a, ASIC1b, ASIC2a, ASIC2b, ASIC3, and ASIC4) (Wemmie et al., 2013). These proteins assemble as homo- or heterotrimers to form channels that are proton-gated, voltage-insensitive, permeable to both Na^+^ and Ca^2+^ and activated by extracellular protons (Waldmann et al., 1997; Waldmann and Lazdunski, 1998). ASIC1a is widely expressed in many areas of the brain, where it is associated with numerous brain functions and disorders, including hippocampally-dependent learning and memory, anxiety, depression, stroke, neurodegeneration, seizure, inflammation, and nerve injury (Chu and Xiong, 2012; Gao et al., 2005; Ortega-Ramírez et al., 2017; Wang et al., 2018; Wemmie et al., 2002; Wemmie et al., 2006). ASIC1a is highly expressed in the amygdala and its ion channel activity is evoked by a reduction in extracellular pH within the physiological range (Taugher et al., 2017; Wemmie et al., 2003). Disruption of ASIC1a affects synaptic transmission and plasticity (Mango and Nisticò, 2019; Soto et al., 2018; Wemmie et al., 2003; Wu et al., 2004). Loss of ASIC1a function also leads to impaired high-frequency electrical stimulation-induced long-term potentiation (LTP) (Chiang et al., 2015; Du et al., 2014; Liu et al., 2016; Wemmie et al., 2002). Moreover, we demonstrated that presynaptic stimulation induces a transient synaptic drop in pH and activates ASIC-like excitable postsynaptic currents (EPSCs) in pyramidal neurons (Du et al., 2014; Kreple et al., 2014), suggesting that protons sufficiently activate postsynaptic ASIC1a in synaptic transmission. In mice, disruption of ASIC1a activity reduces innate fear learning and memory and alters neuronal activity in the fear circuit (Coryell et al., 2007), whereas its overexpression has opposite effects (Wemmie et al., 2004). Also, reducing pH by CO_2_ inhalation or the injection of acid into the amygdala induced aversive behavior response and enhanced aversive memory, and the CO_2_ effects are ASIC1a dependent (Ziemann et al., 2009).

Although our previous data suggested that CO_2_ inhalation affects the aversive memory retrieval and alters the memory lability, we also know that the CO_2_ - affected memory trace is associated with the retrieval cue (Du et al., 2017), inhalation of CO_2_ may reduce pH outside of the amygdala (Dulla et al., 2005; Zandbergen et al., 1989). Thus, whether CO_2_ and ASICs specifically regulate memory trace in retrieval is still unknown and this question is outstanding. In this study, we found that CO_2_ inhalation paired with memory retrieval selectively potentiated memory lability in mice. Furthermore, electrorheological and imaging studies in brain slices support the conclusion that the effects of CO_2_ on memory retrieval are specifically associated with a given memory. Our study proposes that inhaling CO_2_ within the reconsolidation window regulates aversive memory with specificity, providing a unique angle to further study the mechanism by which memory is modulated.

## EXPERIMENTAL PROCEDURES

### Mice

For our experiment, we used both male and female mice between 10-14 months of age. Mice were derived from a congenic C57BL/6 background including wild-type, ASIC1a knock out (ASIC1a^-/-^), and TetTag-c-fos-tTA mice. TetTag-cFos-tTA mice were obtained from Jackson Laboratory and crossed with C57BL/6J mice. Mice carrying the Fos-tTA transgene were selected; Fos-tTA mice have a Fos promoter driving expression of nuclear-localized, two-hour half-life EGFP (shEGFP) (Du et al., 2017; Koffman and Du, 2017; Ramirez et al., 2013). The Fos promoter also drives the expression of tetracycline transactivator (tTA), which bind to the tetracycline-responsive element (TRE) site on an injected recombinant adeno-associated virus, AAV_2/9_-TRE-mCherry virus, resulting in the expression of mCherry (Du et al., 2017; Koffman and Du, 2017). The binding of the tTA to the TRE site is inhibited by doxycycline (DOX). Inhibition of tTA binding prevents target gene expression (Das et al., 2016; Liu et al., 2012; Ramirez et al., 2013).

Both male and female mice age10-14 weeks were randomly selected for the experiment groups. Experimental mice were maintained on a standard 12-hour light-dark cycle and received standard chow and water ad libitum. Animal care and procedures met the National Institutes of Health standards. The University of Tennessee Health Science Center Laboratory Animal Care Unit (Protocol #19-0112) and University of Toledo Institutional Animal Care and Use Committee (Protocol #108791) approved all procedures.

### Aversive conditioning, retrieval, extinction, and memory test

The protocols for each experiment are detailed in the schematics of each figure. All mice were handled by experimenters for 30 minutes on each of 3 days before aversive conditioning. On day 1, mice were habituated to the aversive conditioning chamber (Med Associates Inc.) for 7 minutes. Mice were then exposed to varying conditioning protocols, as described below.

#### Experiment 1: Standard conditioned stimulus (CS) auditory aversive conditioning, retrieval, extinction, and memory tests

On day 1 in a curated environment (context A), the experimental mice were presented with six pure tones (80 dB, 2 kHz, 20 seconds each) paired with 6 foot shocks - one shock at the end of each tone (0.7 mA, 2 seconds). The interval between each tone was 100 seconds. On day 2, the mice were placed into a new environment (context B) and habituated for 4 minutes. Mice then inhaled either unaltered air or air containing 10% CO_2_ for 7 minutes. Five minutes after inhalation of CO_2_ or air began, mice were presented with one 20 second pure tone to retrieve the memory. The mice were then returned to their home cages. 30 minutes later, the mice returned to the retrieval chamber (context B) and underwent two rounds of extinctions. In the first round of extinction, mice were exposed to 20 pure tones with an interval between tones of 100 seconds. Mice were then returned to their home cage. 30 minutes later, the mice went through the extinction protocol again with 20 pure tones. On day 7, the mice were tested to see if their aversive response would recur via spontaneous recovery in context B with 4 pure tones. Thirty minutes after spontaneous recovery, the mice were returned to the original context of the aversive memory, context A, in a recovery protocol with 4 pure tones.

Freezing behavior in mice (the absence of movement beyond respiration) is used as a measure of aversive response. To evaluate the outcomes of freezing behavior in mice, the percentage of time during CS presentation spent in freezing was scored automatically using VideoFreeze software (Med Associates Inc.). In the spontaneous recovery and renewal tests, outcomes of the percentage of time freezing were averaged from each of the 4 CSs.

#### Experiment 2: Two distinct CSs aversive conditioning, retrieval, extinction, and memory tests

This procedure was used to test the specificity of the effects of CO_2_ on memory retrieval. The context settings and parameters are similar to the above standard one CS auditory aversive conditioning. In contrast to experiment 1, the mice were presented with three pure tones (80 dB, 2 kHz, 20 seconds each) that alternated with three white noises (60 dB, 2 kHz, 20 seconds each); all six stimuli were paired with foot shocks. On day 2, the mice inhaled either unaltered air or air containing 10% CO_2_ for 7 minutes. Five minutes after inhalation of CO_2_ or air began, the mice underwent retrieval with one single pure tone followed by one white noise with or without CO_2_. This was followed thirty minutes later by two sections of extinctions with either pure tones or white noises. On day 7, the mice were tested via spontaneous recovery and renewal protocols with 4 pure tones and 4 white noises respectively.

#### Experiment 3: Two distinct CSs aversive conditioning, retrieval, anisomycin, and memory tests

In a series of experiments, the standard extinction procedure was replaced with amygdala infusion of anisomycin (detailed in the surgery procedure below). In brief, the cannula was implanted on the amygdala 4-7 days before the behavioral experiments. Starting on day 1, the mice were subjected to the aversive conditioning described in experiment 2. On day 2, 30 minutes after retrieval, the mice were infused with 62.5 μg/μl anisomycin via the cannula in the lateral nuclei of the amygdala (LA) bilaterally and returned to their home cage (Debiec et al., 2010). On day 7, the mice were tested via spontaneous recovery and renewal as described in experiment 2.

### Surgery and chemical infusion

For the cannula placement procedure, mice were anesthetized with isoflurane through an anesthetic vaporizer, secured to the stereotaxic instrument and a cannula made from a 25-gauge needle was inserted bilaterally into LA (relative to bregma: −1.2 mm anterioposterior; ±3.5 mm mediolateral; −4.3 mm dorsoventral) (Du et al., 2017; Koffman and Du, 2017). Dental cement secured the cannula and bone anchor screw in place. Mice recovered for 4-5 days before any subsequent testing was carried out. A 10 μL Hamilton syringe connected to a 30-gauge injector was inserted 1 mm past the cannula tip to inject anisomycin (diluted in 1 μl artificial cerebrospinal fluid (ACSF), pH 7.3) over 5 sec. The injection sites were mapped post-mortem by sectioning the brain (10 μm coronal) and performing cresyl violet staining.

### Brain slice preparation and patch-clamp recording of amygdala neurons

Ten minutes after the memory retrieval experiment ended, mice were euthanized with overdosed isoflurane and whole brains were dissected into pre-oxygenated (5% CO_2_ and 95% O_2_) ice-cold high sucrose dissection solution containing (in mM): 205 sucrose, 5 KCl, 1.25 NaH_2_PO_4_, 5 MgSO_4_, 26 NaHCO_3_, 1 CaCl_2_, and 25 glucose (Du et al., 2017). A vibratome sliced brains coronally into 300 μm sections that were maintained in normal ACSF containing (in mM): 115 NaCl, 2.5 KCl, 2 CaCl_2_, 1 MgCl_2_, 1.25 NaH_2_PO_4_, 11 glucose, 25 NaHCO_3_ bubbled with 95% O_2_/5% CO_2_, pH 7.35 at 20°C-22°C. Slices were incubated in the ACSF at least 1 hour before recording. For experiments, individual slices were transferred to a submersion-recording chamber and were continuously perfused with the 5% CO_2_/95% O_2_ solution (~3.0 ml/min) at room temperature (20°C - 22°C).

As we described previously (Du et al., 2017), pyramidal neurons in the lateral amygdala were studied using whole-cell patch-clamp recordings. The pipette solution containing (in mM): 135 KSO_3_CH_3_, 5 NaCl, 10 HEPES, 4 MgATP, 0.3 Na3GTP, 0.5 K-EGTA (mOsm=290, adjusted to pH 7.25 with KOH). The pipette resistance (measured in the bath solution) was 3-5 MΩ. High-resistance (>1 GΩ) seals were formed in voltage-or 20 white noiseclamp mode. Picrotoxin (100 μM) was added to the ACSF throughout the recordings to yield excitatory responses. In AMPAR current rectification experiments, we applied D-APV (100 μM) to block NMDAR-conducted EPSCs. The peak amplitude of ESPCs was measured to determine current rectification. The amplitude was measured ranging from −80 mV to +60 mV in 20 mV steps. The peak amplitude of EPSCs at −80 mV and +60 mV was measured for the rectification index. In EPSC ratio experiments, neurons were measured at −80 mV to record AMPAR-EPSCs and were measured at +60 mV to record NMDAR-EPSCs. To determine the AMPAR-to-NMDAR ratio, we measured the peak amplitude of ESPCs at −80 mV as AMPAR-currents, and peak amplitude of EPSCs at +60 mV at 70 ms as NMDAR-currents after onset. Data were acquired at 10 kHz using Multiclamp 700B and pClamp 10.1. The mEPSCs events (>5pA) were analyzed in Clampfit 10.1. The decay time (τ) of mEPSCs was fitted to an exponential using Clampfit 10.1.

### Immunohistochemistry and cell counting

The AAV-TRE-mCherry plasmid was obtained from the laboratory of Dr. Susumu Tonegawa (Liu et al., 2012; Ramirez et al., 2013), and was used to produce AAV_2/9_ by the University of Iowa Gene Transfer Vector Core. For one week leading up to virus microinjection, TetTag Fos-tTA mice were fed with food containing 40 mg/kg DOX. We used a 10 μl Hamilton microsyringe and a WPI microsyringe pump to inject virus (0.5 μl of 1.45E+12 viral genomes/ml of AAV_2/9_-TRE-mCherry) bilaterally into the amygdala (relative to bregma: −1.2 mm anterioposterior; ±3.5 mm mediolateral; −4.3 mm dorsoventral), as described previously (Du et al., 2017; Koffman and Du, 2017). For a two-week window between surgery and behavior training, mice were housed and fed with a DOX-containing diet. The DOX-containing diet was ceased twenty-four hours before aversive conditioning began on day one (replaced by a regular diet), then immediately restarted afterward. Thirty minutes after retrieval on day two, the mice were euthanized according to protocol. We used transcardial perfusion with 4% paraformaldehyde (PFA) to fix whole brains, followed by continued fixation in 4% PFA at 4°C for 24 hours (Wright et al., 2020). Following perfusion, we used a vibratome (Leica VT-1000S) to dissect 50 μm amygdala coronal slices, which were collected in ice-cold PBS. To complete immunostaining, slices were placed in Superblock solution (Thermo Fisher Scientific) plus 0.2% Triton X-100 for 1 hour and incubated with primary antibodies (1:1000 dilution) at 4°C for 24 hours (Du et al., 2017). Primary antibodies we used include: rabbit polyclonal IgG anti-RFP (Rockland Cat# 600-401-379); chicken IgY anti-GFP (Thermo Fisher Scientific Cat# A10262) and mouse anti-NeuN (Millipore Cat# MAB377X) (Liu et al., 2012; Ramirez et al., 2013). We then washed and incubated slices for one hour with secondary antibodies (Alexa Fluor 488 goat anti-chicken IgG (H+L) (Molecular Probes Cat# A-11039); Alexa Fluor 568 goat anti-rabbit IgG (H+L) (Molecular Probes Cat# A-21429); Alexa Fluor 647 goat anti-mouse IgG (H+L) (Thermo Fisher Scientific Cat# A-21235), 1:200 dilution). VectaShield H-1500 (Vector Laboratories Cat# H-1500) was used to mount slices, while confocal microscopy was used to view the slices. We used ImageJ software to analyze dendritic spine morphology. Thin, mushroom and stubby spines were categorized based on the following parameters: 1) mushroom spines: head-to-neck diameter ratio >1.1:1 and spine head diameter >0.35 μm; 2); thin spines: head-to-neck diameter ratio >1.1:1 and spine head diameter >0.35 μm or spine head-to-neck diameter ratios <1.1:1 and spine length-to-neck diameter > 2.5 μm; 3); stubby spines: spine head-to-neck diameter ratios <1.1:1 and spine length-to-neck diameter ≤ 2.5 μm (Kreple et al., 2014; Wright et al., 2020).

### Statistical analysis

One-way ANOVA and Tukey’s post-hoc multiple comparison tests were used for statistical comparison of groups. An unpaired Student’s t-test was used to compare results between two groups. P<0.05 was considered statistically significant, and we did not exclude potential outliers from our data except the ones did not receive successful aversive conditioning. The graphing and statistical analysis software Graphpad Prism 8 was used to analyze statistical data, which was presented as means ± SEM. Sample sizes (n) are indicated in the figure legends, and data are reported as biological replicates (data from different mice, different brain slices). Each group contained tissues pooled from 4-5 mice. Due to variable behavior within groups, we used sample sizes of 10-16 mice per experimental group as we previously described in earlier experiments (Du et al., 2017). In behavioral studies, we typically studied groups with four randomly assigned animals per group, as our recording equipment allowed us to record four separate animal cages simultaneously. The experiments were repeated with another set of four animals until we reached the target number of experimental mice per group. Experimentation groups were repeated in this manner so that each animal had the same controlled environment-the same time of day and with similar handling, habituation, and processes.

## RESULTS

Our recent studies suggested that CO_2_ inhalation throughout memory retrieval enhances the lability of the memory and boosts the efficiency of the memory erasure (Du et al., 2017). To further test whether the effects of CO_2_ on memory retrieval are synapse-specific, we designed a series of unique experiments by which we were able to generate two distinct auditory aversive memories and identify the specificity of CO_2_ effects on each of them.

### CO_2_ selectively enhances the lability of auditory aversive memory in the amygdala

Previous studies have described an aversive conditioning paradigm in which memory can be selectively reactivated and reconsolidated, suggesting synapse-specific reconsolidation of distinct aversive memories in the amygdala (Debiec et al., 2010; Doyere et al., 2007). We followed this paradigm albeit with modifications **(Fig.1A, Fig. s1A)**. On day 1, we trained the animals with two distinct conditioned stimuli: three pure tones and three white noises paired with one foot-shock per stimuli as aversive unconditioned stimuli (US) (see the detailed description in Materials and Methods). We evaluated the outputs of the aversive conditioning through the percentage of the freezing time within the time of CSs. The freezing was significantly increased after each of the three conditioned stimuli, indicating the mice were trained sufficiently under the designed condition **(Fig.1B, Fig. s1B)**.

**Figure 1:**
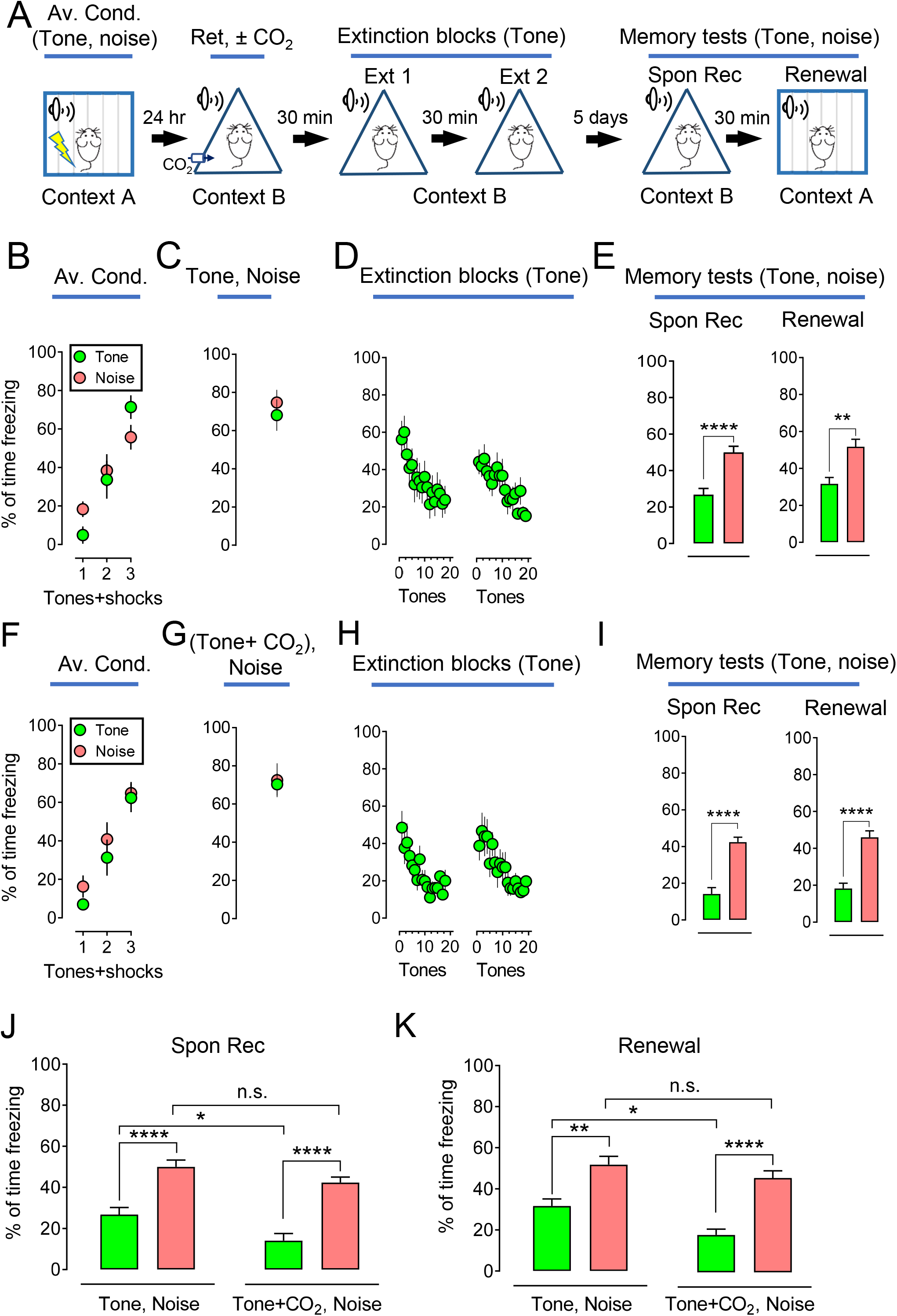
CO_2_ inhalation during a selective memory retrieval potentiates the effect of the extinction. **(A)** Representative schematic of protocol for the aversive conditioning (Av. Cond.), memory retrieval (Ret), extinction (Ext), and memory test-spontaneous recovery (Spon Rec) and renewal. On day1, the mice were subjected to 3 pure tones and 3 white noises, paired with 6 foot shocks in context A. One day after, the mice were placed in context B and were subjected to both tone and noise as retrieval events. 30 mins after retrieval, the mice were treated with 2 blocks of extinctions with a pure tone as the CS. On day 7, spontaneous recovery and renewal were tested individually in context B and then context A. 4 pure tones and 4 white noises were presented as CSs during each memory testing. **(B-E)** Data are presented by the percentage of freezing time during the CSs (tone and noise) in aversive conditioning **(B)**, retrievals (tone and noise) **(C)**, two sections of extinction with tone **(D)**, memory test of spontaneous recovery (Spon Rec) and renewal with tone and noise **(E)**. **(F-I)** Data are presented by the percentage of freezing time during the CSs (tone and noise) in aversive conditioning **(F)**, retrievals (tone plus CO_2_ inhalation and noise) **(G)**, two sections of extinction with tone **(H)**, memory test of spontaneous recovery (Spon Rec) and renewal with tone and noise **(I)**. **(J-K),** comparison data based on spontaneous recovery and renewal respectively from panels **E** and **I**. Data are mean ± SEM. n = 10-12 mice in each group. ‘n.s.’ indicates not statistically significant. * indicates p<0.05, ** indicates p<0.01, *** indicates p<0.001, **** indicates p<0.0001,by unpaired Student’s t-test and one-way ANOVA with Tukey’s post-hoc multiple comparisons. See also Figure S1 and Figure S2.

On day 2, the animals were placed into a new context (context B) and presented with one pure tone followed by a single noise (or vice versa) to retrieve the memory **(Fig. 1A, C, Fig. s1A, C)**. The animals were then returned to their home cages. 30 minutes later, all mice underwent two blocks of extinctions in context B, each extinction contains 20 pure tones **(Fig.1D)** or 20 white noise **(Fig. s1D).** At the end of the extinction procedure, the freezing dropped down to a low level **(Fig. 1D, Fig. s1D)**, suggesting that the extinction procedure was sufficient to suppress the memory. Five days later, the mice underwent spontaneous recovery (context B) and renewal (context A) respectively. Four tones and four noises were presented throughout the memory test **(Fig.1A, Fig. s1A)**. The group that underwent extinction with a specific CS showed specificity in which freezing response was lowered after extinction **(Fig.1E, Fig. s1E)**. For example, when the pure tone was presented during extinction, freezing in the pure tone group was lower than freezing in the noise group **(Fig.1E)**, and vice versa **(Fig. s1E)**. When retrieval was paired with 10% CO_2_ inhalation, memory erasure effects were enhanced **(Fig.1 F-I, Fig. s1F-I)**, and there are statistical significances between the groups with or without CO_2_ in both spontaneous recovery and renewal (green columns, **Fig.1 J-K** and red columns, **Fig. s1J-K**). To further evaluate the specificity of the effects of reconsolidation on memory modifications, we designed another retrieval protocol in which we presented pure tone and white noise, either of them paired with 10% CO_2_ inhalation, followed by an unrelated extinction of CS, white noise, or pure tone respectively **(Fig. s2)**. Consistent with our expectation, CO_2_ did not boost the effects on the retrieved memory in the absence of paired extinction, compared to the data in Fig. 1 and Fig. S1 **(Fig. s2B-I)**. When compared to the data in **Fig. 1F-I** and **Fig. s1F-I**, we found the application of 10% CO_2_ to the retrieval event failed to enhance the outcome after extinction, indicating a specificity of the CO_2_ effects. In all, our data suggest that memory encoded in the amygdala can be distinct, and the effects of CO_2_ on memory are specific.

To focus on testing the specific effects of CO_2_ on retrieval, we replaced the extinction procedure with an injection of a protein synthesis inhibitor, anisomycin, to obliterate the aversive memory **(Fig. 2A)**. Anisomycin, when injected bilaterally into the amygdala after retrieval, causes memory erasure compared to the saline injection group (Debiec et al., 2010). We conditioned the mice and retrieve the memory with pure tones **(Fig. 2B-C)**, followed by anisomycin/saline injection **(Fig. 2D)**. Our experiments show that anisomycin disrupts the memory during reconsolidation **(Fig. 2E)**, and that memory retrieval is required for memory erasure with anisomycin injection **(Fig. s3A-E)**. Consistent with the extinction results in Fig. 1, when retrieval was paired with 10% CO_2_ inhalation, we found anisomycin reduced more aversive response-further confirming that CO_2_ enhances memory lability specifically **(Fig. 2F-I)**. To exclude the possibility that anisomycin associates with one CS other the other, as rigorous controls, we conditioned the mice with both pure tone and white noise and carried out memory retrieval with both CSs and found anisomycin has equal effects on memory in both tone and noise groups **(Fig. s3F-M)**. When 10% CO_2_ was applied while the CSs were presented, the retrieval group paired with CO_2_ showed less freezing, regardless of the type of CSs (pure tone or white noise) **(Fig.s4 B-I)**. As rigorous controls, we applied CO_2_ for both retrieval events together and anisomycin decreased the freezing level in both groups, suggesting that CO_2_ had equal effects on both tone and noise **(Fig. s4 J-Q)**. As another control, saline injection following retrieval and CO_2_ did not cause memory erasure, indicating that anisomycin was necessary to disrupt the reactivated memory **(Fig. s5)**. Talking together, our data demonstrate that the effects of CO_2_ are specific to a distinct memory that is activated by a specific CS.

**Figure 2:**
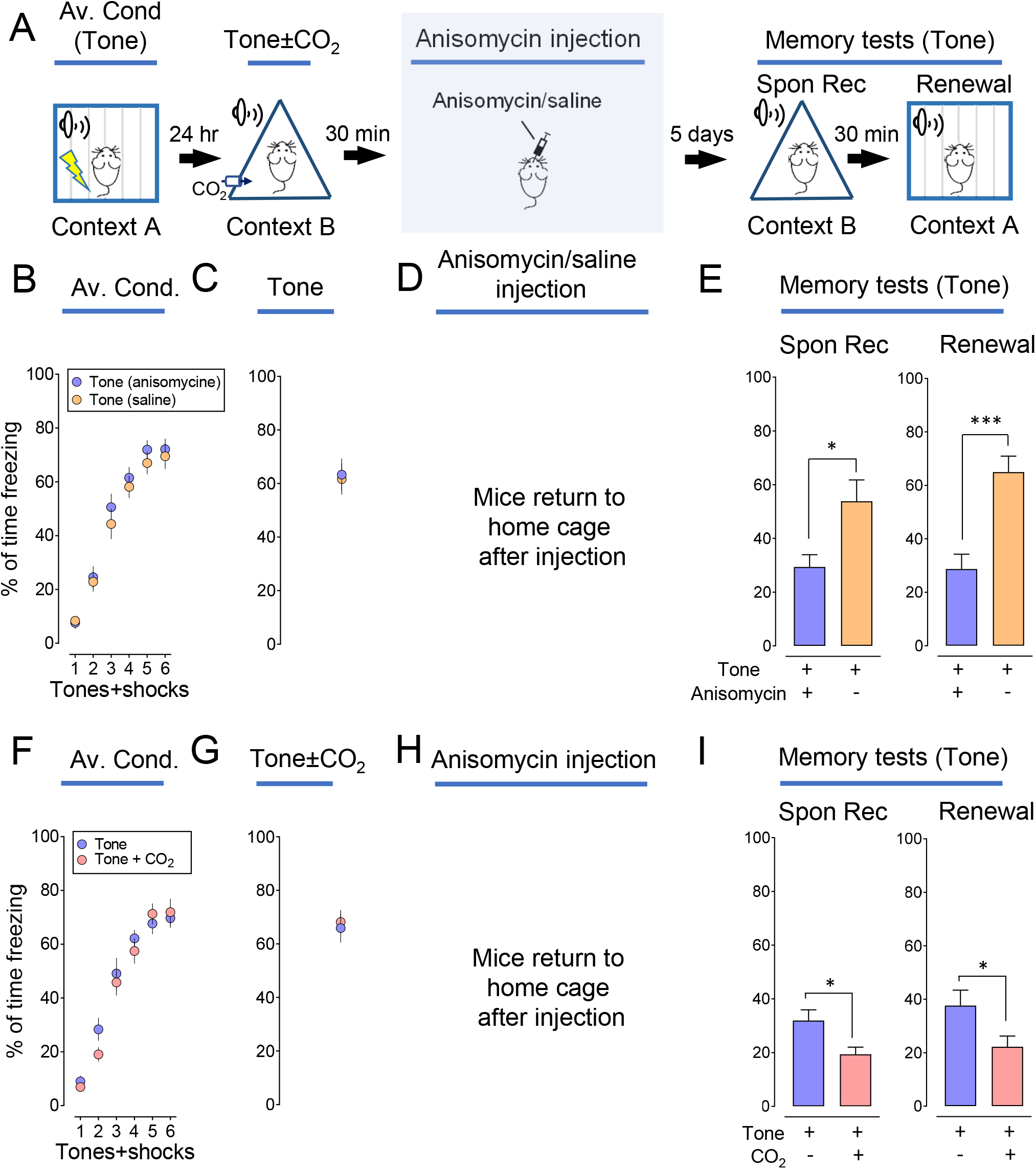
CO_2_ inhalation during memory retrieval potentiates the effect of the anisomycin. **(A)** Representative schematic of the protocol for aversive conditioning, memory retrieval, anisomycin injection, and memory test (spontaneous recovery and renewal). On day1, the mice were subjected to 6 pure tones, paired with 6 foot shocks in context A. One day after CS, the mice were placed in context B and were subjected to one single tone as a retrieval event with or without CO_2_ inhalation. 30 mins after retrieval, the mice were then infused with 62.5 μg/μl anisomycin or saline and then returned to their home cage. On day 7, spontaneous recovery and renewal were tested individually in context B and then context A. 4 pure tones were presented as CSs during each memory testing. **(B-E)** Data are presented by the percentage of freezing time during the tone presentation in aversive conditioning **(B)**, retrieval (tone) **(C)**, saline or anisomycin infusion in the amygdala **(D)**, spontaneous recovery (Spon Rec), and renewal test with tones **(E)**. **(F-I)** Data are presented by the percentage of freezing time during the tone presentation in aversive conditioning **(F)**, retrieval (tone) with or without CO_2_ **(G)**, anisomycin infusion **(H)**, spontaneous recovery (Spon Rec), and renewal test with tones **(I)**. Data are mean ± SEM. n = 8-10 mice in each group. * indicates p<0.05, *** indicates p<0.001, by unpaired Student’s t-test. See also Fig. s3, s4, s5

### The specific effects of CO_2_ on memory lability is ASIC-dependent

We have previously found the effects of CO_2_ on memory retrieval to be ASIC dependent (Du et al., 2017). However, it is still unknown if CO_2_ application to a specific memory trace affects an ASIC-dependent mechanism. To answer this question, we first performed distinct aversive conditioning in ASIC1a^-/-^mice with three pure tones and white noises on day 1 **(Fig. 3A)**, followed by a pure tone and white noise for retrieval on day 2. 30 minutes post-retrieval, we performed extinctions with pure tones. Five days later, we tested spontaneous recovery and renewals with 4 pure tones and white noises. Similar to the response we saw in WT mice, the freezing level in the pure tone group of ASIC1a^-/-^ mice were less than that in the white noise group (spontaneous recovery, 46% decrease; renewal, 47.5% decrease) **(Fig. 3B-E)**. When 10% CO_2_ inhalation was paired with pure tone in retrieval, we found that CO_2_ did not have additional effects on the memory with the specific CS in ASIC1a^-/-^ mice (spontaneous recovery, 43.4% decrease; renewal, 45.6% decrease) **(Fig. 3F-I)**, and there are no statistical significances between the groups with or without CO_2_ in both spontaneous recovery and renewal (green columns, **Fig. 3J, K**). We had hypothesized that the effects of CO_2_ on memory retrieval would be ASIC dependent and our data supported this prediction. We then replaced the extinction procedure with anisomycin, to obliterate the aversive memory **(Fig. 4A)**. The ASIC1a^-/-^ mice were conditioned with pure tones **(Fig. 4B)** and followed by a single tone as retrieval **(Fig. 4C)**. We then apply anisomycin infusions **(Fig. 4D)**and test the memory 5 days later **(Fig. 4E)**. Anisomycin dramatically reduced freezing in memory tests that followed, whereas pairing with CO_2_ in the CS did not cause an additional reduction in the ASIC1a^-/-^ mice, suggesting an ASIC dependency **(Fig. 4E)**. We also presented two distinct CSs (pure tone and white noise) during conditioning and then retrieval, with or without 10% CO_2_, followed by anisomycin infusions in the ASIC1a^-/-^ mice **(Fig. 4F)**. Anisomycin dramatically reduced freezing in memory tests that followed, whereas pairing with CO_2_ in either CSs (pure tone or white noise) did not cause an additional reduction in the ASIC1a^-/-^ mice, suggesting an ASIC dependency **(Fig. 4G-N)**.

**Figure 3:**
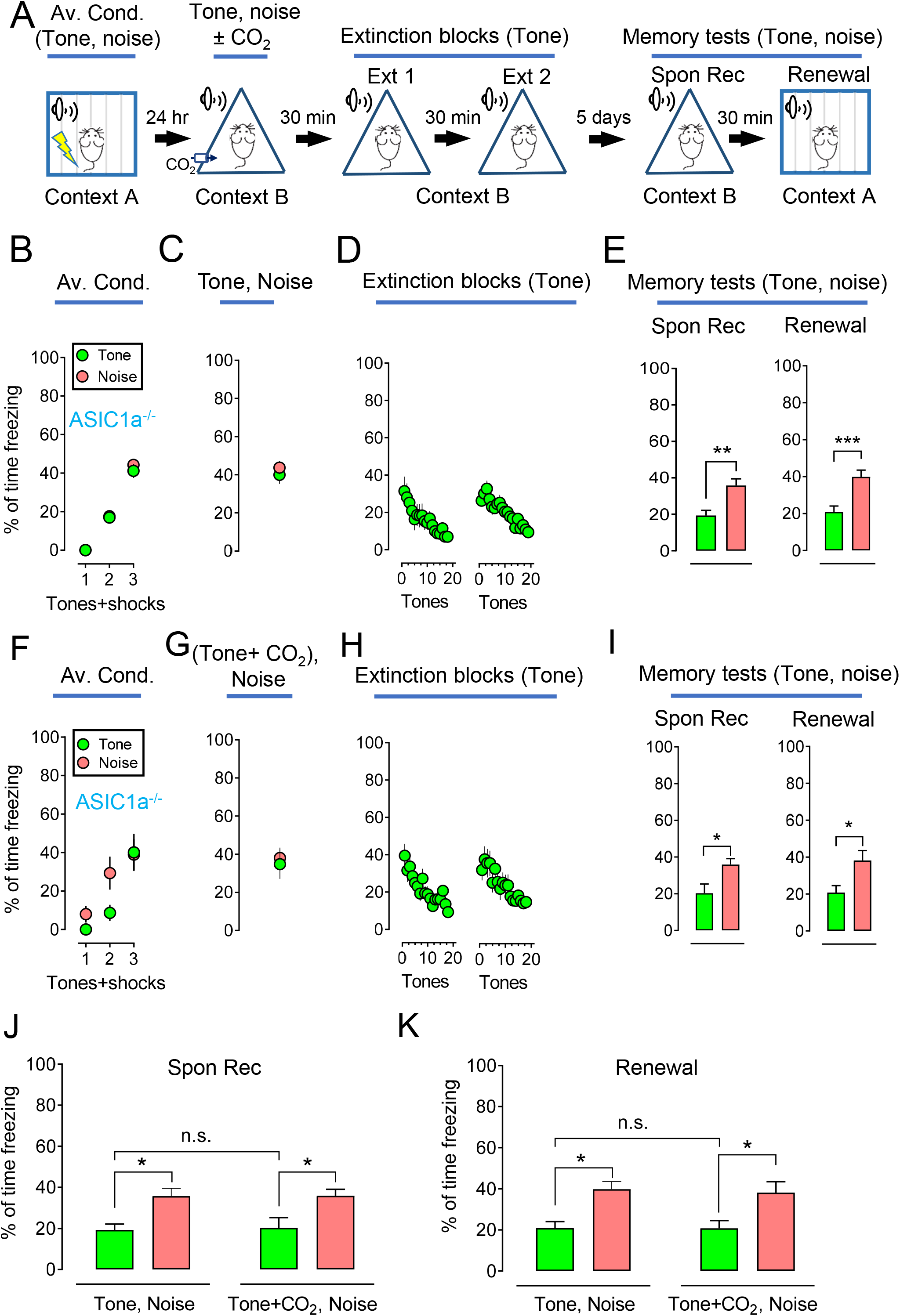
The effect of CO_2_ inhalation on selective memory retrieval is ASIC1a-dependent. **(A)** Representative schematic of protocol for the aversive conditioning, memory retrieval, extinction, and memory test (spontaneous recovery and renewal) in ASIC1a^-/-^ mice. On day1, the mice were subjected to 3 pure tones and 3 white noises, paired with 6 foot shocks in context A. One day later, the mice were placed in context B and subjected to both tone and noise as retrieval events. 30 mins after retrieval, the mice were treated with 2 blocks of extinctions with a pure tone as the CS. On day 7, spontaneous recovery and renewal were tested individually in context B and then context A. 4 pure tones and 4 white noises were presented as CSs during each memory testing. **(B-E)** Data in ASIC1a^-/-^ mice are presented by the percentage of freezing time during the CSs (tone and noise) in aversive conditioning **(B)**, retrievals (tone and noise) **(C)**, two sections of extinction with tone **(D)**, memory test of spontaneous recovery (Spon Rec) and renewal with tone and noise **(E)**. **(F-I)** Data in ASIC1a^-/-^ mice in aversive conditioning **(F)**, retrievals (pure tone plus 10% CO_2_ inhalation and white noise) **(G)**, two sections of extinction with pure tone **(H)**, memory test of spontaneous recovery (Spon Rec) and renewal with tone and noise **(I)**. **(J-K),** comparison data based on spontaneous recovery and renewal respectively from panels **E** and **I**. Data are mean ± SEM. n = 12-16 mice in each group. * indicates p<0.05, ** indicates p<0.01, *** indicates p<0.001, by unpaired Student’s t-test and one-way ANOVA with Tukey’s post-hoc multiple comparisons.

**Figure 4:**
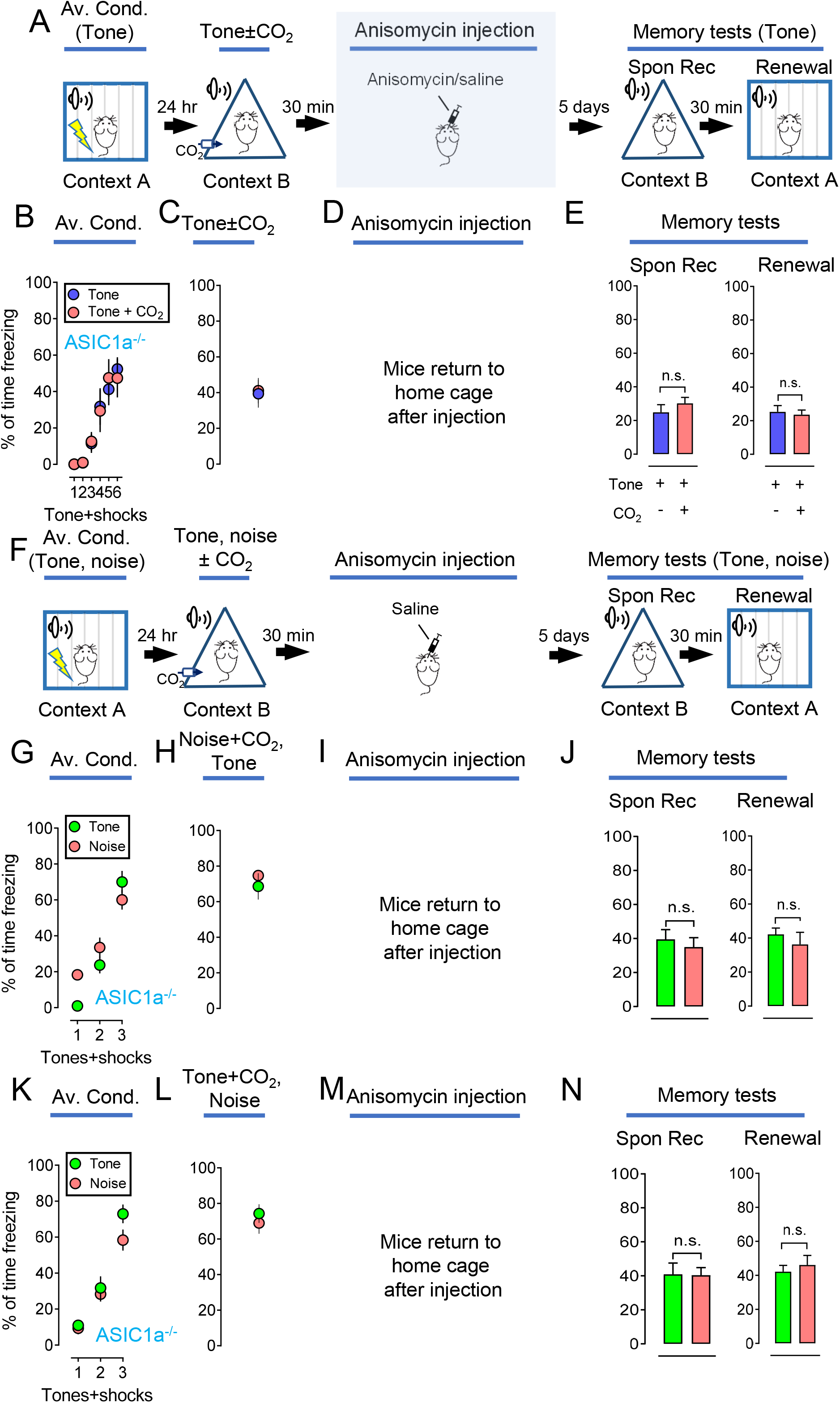
The CO_2_ potentiated anisomycin effects are ASIC1a-dependent. **(A)** Representative schematic of protocol for the aversive conditioning, memory retrieval, anisomycin injection, and memory test (spontaneous recovery and renewal). On day 1, the mice were subjected to 6 pure tones, paired with 6 foot shocks in context A. One day after, the mice were placed in context B and subjected to one single tone as a retrieval event with or without CO_2_ inhalation. 30 mins after retrieval, the mice were infused with 62.5 μg/μl anisomycin or saline and then returned to their home cage. On day 7, spontaneous recovery and renewal were tested individually in context B and then context A. 4 pure tones were presented as CSs during each memory testing. **(B-E)** Data in ASIC1a^-/-^ mice are presented by the percentage of freezing time during the tone presentation in aversive conditioning **(B)**, retrieval (tone) with or without CO_2_ **(C)**, anisomycin infusion in the amygdala **(D)**, spontaneous recovery (Spon Rec) and renewal test with tones **(E)**. **(F)** Representative schematic of protocol for the aversive conditioning, memory retrieval, extinction, and memory test (spontaneous recovery and renewal). On day 1, ASIC1a^-/-^ mice were subjected to 3 pure tones and 3 white noises, paired with 6 foot shocks in context A. One day later, the mice were placed in context B and subjected to both tone and noise as retrieval events. 30 mins after retrieval, the mice were infused with anisomycin or saline and returned to their home cage. On day 7, spontaneous recovery and renewal were tested individually in context B and then context A. 4 pure tones and 4 white noises were presented as CSs during each memory testing. **(G-J)** Data are presented by the percentage of freezing time during the tone presentation in aversive conditioning **(G)**, retrieval (noise plus CO_2_ inhalation, then tone) **(H)**, anisomycin infusion in the amygdala **(I)**, spontaneous recovery (Spon Rec), and renewal test with tones **(J)**. **(K-N)** Data are presented by the percentage of freezing time during the tone presentation in aversive conditioning **(K)**, retrieval (tone) with or without CO_2_ **(L)**, anisomycin infusion in the amygdala **(M)**, spontaneous recovery (Spon Rec), and renewal test with tones **(N)**. Data are mean ± SEM. n = 12-16 mice in each group. ‘n.s.’ indicates not statistically significant by unpaired Student’s t-test between groups.

To provide evidence that an acute ASIC1a blockage was able to eliminate the effects of CO_2_, we injected 100nM PcTX-1 into the lateral amygdala bilaterally 1 hour before the application of CO_2_ to the retrieval **(Fig. 5A)**. Our data suggest that compared to the saline injection group **(Fig. 5 B-E)**, inhibiting ASIC1a by PcTX-1 significantly reduced the CO_2_ effects on memory retrieval **(Fig. 5 F-I)**, statistical analysis in the spontaneous recovery and renewal groups supported this conclusion (green columns, **Fig. 5J, K**). This pattern of findings suggests that the effects of CO_2_ on specific memory traces are ASIC dependent.

**Figure 5:**
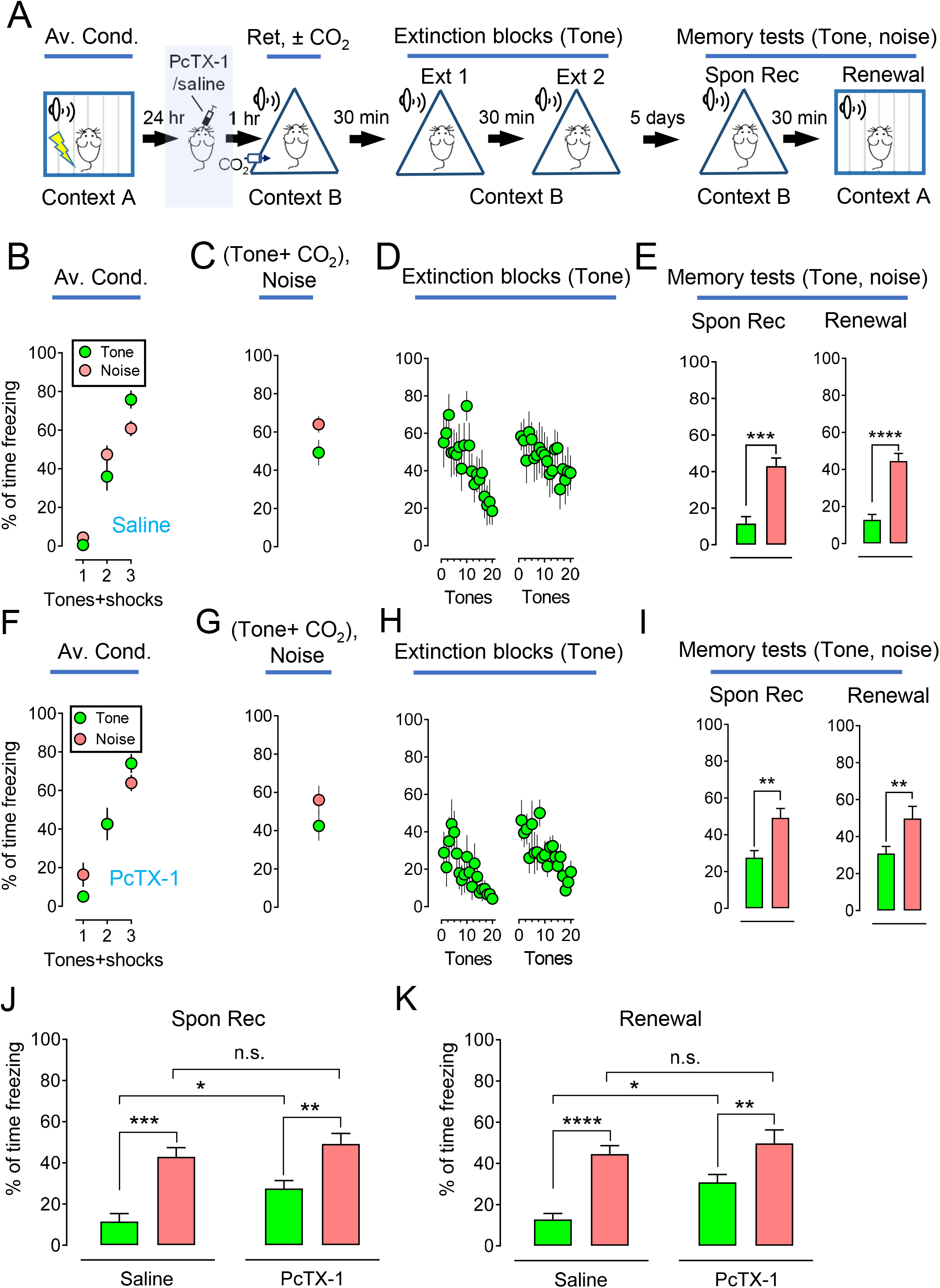
Blockage of ASIC1a in the amygdala reduces the CO_2_ effects on selective memory retrieval. **(A)** Representative schematic of protocol for the aversive conditioning, PcTX-1 injection, memory retrieval, extinction, and memory test (spontaneous recovery and renewal). On day1, the mice were subjected to 3 pure tones and 3 white noises, paired with 6 foot shocks in context A. One day later, the mice were injected with 100nM PcTX-1 or saline, then the mice were placed in context B and subjected to both tone and noise as retrieval events with or without CO_2_. 30 mins after retrieval, the mice were treated with 2 blocks of extinctions with a pure tone as the CS. On day 7, spontaneous recovery and renewal were tested individually in context B and then context A. 4 pure tones and 4 white noises were presented as CSs during each memory testing. **(B-E)** Data are presented by the percentage of freezing time during the CSs (tone and noise) in aversive conditioning **(B)**, retrievals (tone plus CO_2_ inhalation and noise) after saline injection in the amygdala **(C)**, two sections of extinction with tone **(D)**, memory test of spontaneous recovery (Spon Rec) and renewal with tone and noise **(E)**. **(F-I)** Data are presented by the percentage of freezing time during the CSs (tone and noise) in aversive conditioning **(F)**, retrievals (tone plus CO_2_ inhalation and noise) after PcTx-1 injection in the amygdala **(G)**, two sections of extinction with tone **(H)**, memory test of spontaneous recovery (Spon Rec) and renewal with tone and noise **(I)**. **(J-K),** comparison data based on spontaneous recovery and renewal respectively from panels **E** and **I**. Data are mean ± SEM. n = 12 mice in each group. ‘n.s.’ indicates not statistically significant. * indicates p<0.05, ** indicates p<0.01, *** indicates p<0.001, **** indicates p<0.0001, by unpaired Student’s t-test and one-way ANOVA with Tukey’s post-hoc multiple comparisons.

### Activation of ASICs through CO_2_ inhalation alters reconsolidation of distinct memory through alteration of AMPARs

AMPARs are glutamatergic receptors that have crucial roles in modulating memory retrieval and destabilization (Auber et al., 2013; Chan et al., 2010; Clem and Huganir, 2010; Monfils et al., 2009; Quirk et al., 2010). Previous studies suggest that the exchange of Ca^2+^-impermeable AMPARs (CI-AMPARs) for Ca^2+^-permeable AMPARs (CP-AMPARs) occurs after retrieval (Clem and Huganir, 2010; Hong et al., 2013). 10% CO_2_ inhalation paired with retrieval induces a stronger current rectification of AMPARs (the signature of CP-AMPARs) than in the retrieval alone group, indicating a greater exchange of AMPARs (Du et al., 2017). Interestingly, no further enhancement was observed in the ASIC1a^-/-^ brain slices, indicating that the effect of CO_2_ inhalation on AMPAR exchange is ASIC dependent (Du et al., 2017). To further study whether CO_2_ specifically alters the AMPARs exchange in retrieval, we designed a unique experiment to separate the aversive conditioning and retrieval and measure the rectification of AMPARs **(Fig. 6A)**. To study this, we conditioned the mice with 6 pure tones on day 1 **(Fig. 6B, left)**. On day 2, the mice were divided into 4 groups based on retrieval conditions - the first group received pure tone only; the second group-pure tone plus 10% CO_2_ inhalation; the third group - white noise only; and the fourth group received white noise+10% CO_2_ inhalation **(Fig. 6B, right)**. Ten minutes after retrieval, we dissected brain slices and AMPAR current was recorded in the pyramidal neurons in the lateral amygdala through stimulation of thalamic inputs **(Fig. 6A)**. Rectification, a signature of CP-AMPARs, was compared among all groups. Consistent with earlier reports, pure tone retrieval increased current rectification (Clem and Huganir, 2010; Hong et al., 2013) and CO_2_ paired with pure tone retrieval caused stronger rectification **(Fig. 6C)**. However, when white noise was presented as the retrieval event, both white noise and white noise plus CO_2_ failed to cause a significant rectification compared to the pure tone group **(Fig. 6C)**. This data supports our prediction that CO_2_ was associated with a specific memory trace that was reactivated. To control for the possible effects of the order of presentation of CSs, we switched over the pure tone and white noise in the aversive conditioning and retrieval. Similar results were observed, confirming the effects of CO_2_ were not artificial **(Fig. 6D, E)**.

**Figure 6:**
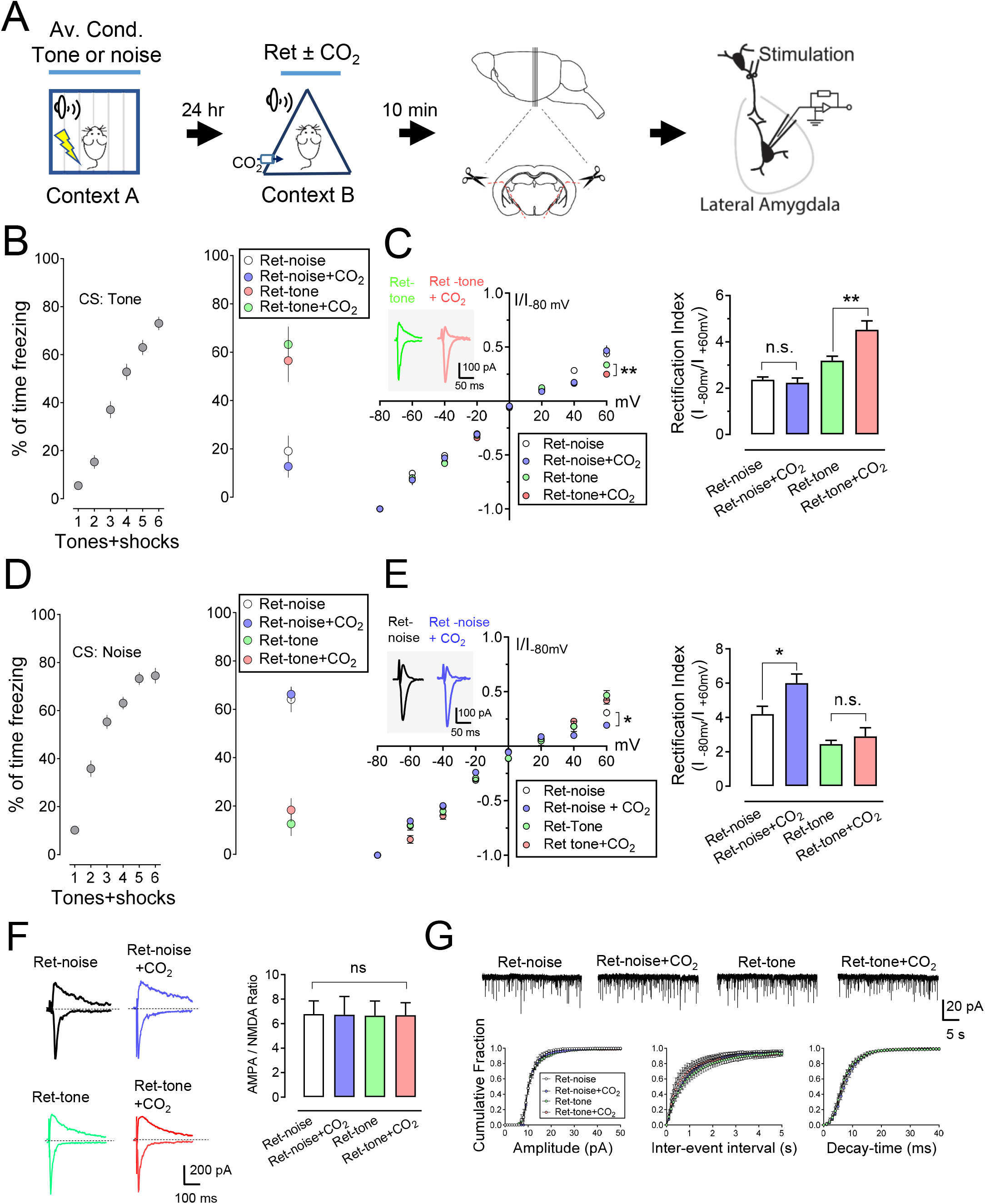
CO_2_ inhalation during a selective memory retrieval enhances the retrieval dependent AMPAR current rectification. **(A)** Representative schematic of the protocol. On day 1, the animal underwent 6 CSs (tones or noses), paired with 6-foot shocks in context A. On day 2, the mice were divided into 4 groups for the retrieval-pure tone only; pure tone plus 10% CO_2_ inhalation; white noise only; white noise+ 10% CO_2_ inhalation. Ten minutes after retrieval, the brain slices were dissected and AMPAR current was recorded in the pyramidal neurons in the lateral amygdala through stimulation of thalamic input. **(B-E)** Mice underwent 6 pure tones **(B)** or 6 noises **(D)** in aversive conditioning, Data are presented by the percentage of freezing time during the tone presentation in aversive conditioning, retrieval (noise plus CO_2_ inhalation, then tone). **(C, E)** Left, AMPARs current-voltage relationships in the recorded neurons. Insets show an example of the AMPAR-EPSCs in −80mV and + 60mV. Right, AMPAR rectification index (I-80 mV / I+60 mV). Data are mean±SEM. n = 20-26 for each group. **(F)** Left, examples of EPSC recordings of AMPAR-EPSCs (−80mV) and NMDAR-EPSCs (+60mV). Right, AMPAR/NMDAR EPSC ratios. Current amplitudes were measured 70 ms after onset. n = 20-26 for each group. **(G)** Miniature EPSCs recordings from the neurons after retrieval. Upper, representative mEPSC traces from different groups. Lower, cumulative distributions of mEPSC amplitudes, inter-event intervals, and decay-times. n = 25-40 for each group. Data are mean ± SEM. ‘n.s.’ indicates not statistically significant. * indicates p<0.05, ** indicates p<0.01, by ANOVA with Tukey’s post hoc multiple comparisons.

We then asked whether synaptic strength changed with the application of an unrelated retrieval CS and CO_2_. The ratio of AMPAR-EPSCs to NMDAR-EPSCs might represent the strength of the synapse (Rao and Finkbeiner, 2007). Previous studies reported that the AMPAR/ NMDAR-EPSC ratio increased after aversive conditioning whereas retrieval did not potentiate further increase, suggesting that memory retrieval did not alter the synaptic strength (Clem and Huganir, 2010; Hong et al., 2013). Our previous studies also indicated that CO_2_ inhalation during memory retrieval did not strengthen the synapse in the amygdala (Du et al., 2017). We further tested whether the pairing of CO_2_ inhalation with the specific retrieval CS influenced the strength of a synapse. Currents were recorded at −80mV for AMPAR-EPSCs and +60mV for NMDAR-EPSCs. Our data suggest that retrieval plus CO_2_ inhalation did not change the AMPAR/ NMDAR-EPSCs ratio **(Fig. 6F)**. Moreover, the characteristics of miniature EPSCs (mEPSCs), including amplitude, frequency, and decay times, were not altered **(Fig. 6G)**. In addition, the pairing of CO_2_ with another unrelated CS in retrieval did not change the strength of the synapse **(Fig. 6F, G)**. This data suggests that the effects of CO_2_ inhalation on specific memory retrieval enhance the destabilization of the synapse without changing synaptic strength.

### The effects of CO_2_ inhalation on distinct memory trace

Our previous studies indicate that CO_2_ enhances memory trace that is associated with aversive conditioning (Du et al., 2017). In this experiment, we examined the mechanism behind the specificity of CO_2_ effects on memory traces. Using the TetTag-c-fos driven-GFP mouse model, neurons in the amygdala involved in memory trace after aversive conditioning can be labeled with a long-lasting mCherry fluorescent protein and the neuron in the retrieval trace can be labeled with a 2-hour short half-life nuclear-localized EGFP (shEGFP) **(Fig. 7A, B)** (see the details in Materials and Methods) (Du et al., 2017; Koffman and Du, 2017). The overlapped labeling (yellow) represents the neurons in the same memory trace of aversive and retrieval (Du et al., 2017; Liu et al., 2012; Ramirez et al., 2013). In this experiment, the mice were first conditioned with pure tone, activating the associated neurons that were labeled with mCherry **(Fig. 7C)**. Right after aversive conditioning, the mice were immediately fed with a DOX diet thereby preventing further mCherry labeling. On day 2, the mice were divided into two groups-one group of mice were presented with a single pure tone to retrieve the memory, another was presented with white noise. A temporary, 2-hour half-life, shEGFP was labeled after the retrieval. Thirty minutes after the retrieval event, we sliced the amygdala and imaged shEGFP-and mCherry-positive cells **(Fig. 7D)**. Compared to pure tone aversive conditioning/pure tone retrieval group (same memory trace of aversive conditioning and retrieval), inhalation of CO_2_ in the pure tone aversive conditioning/white noise retrieval group did not result in an increase of neurons positive for expression of both mCherry-positive and shEGFP-positive neurons (overlapped labeling, yellow) **(Fig. 7E)**. Control experiments to identify the efficiency of the aversive conditioning on the expression of mCherry on the cells were performed **(Fig. 7F)**. These findings indicate that CO_2_, when paired with retrieval, only enhances the memory trace that has been reactivated; CO_2_ paired with unrelated retrieval cue does not affect the original memory trace. These findings suggest a specific effect of CO_2_ on the memory trace.

**Figure 7:**
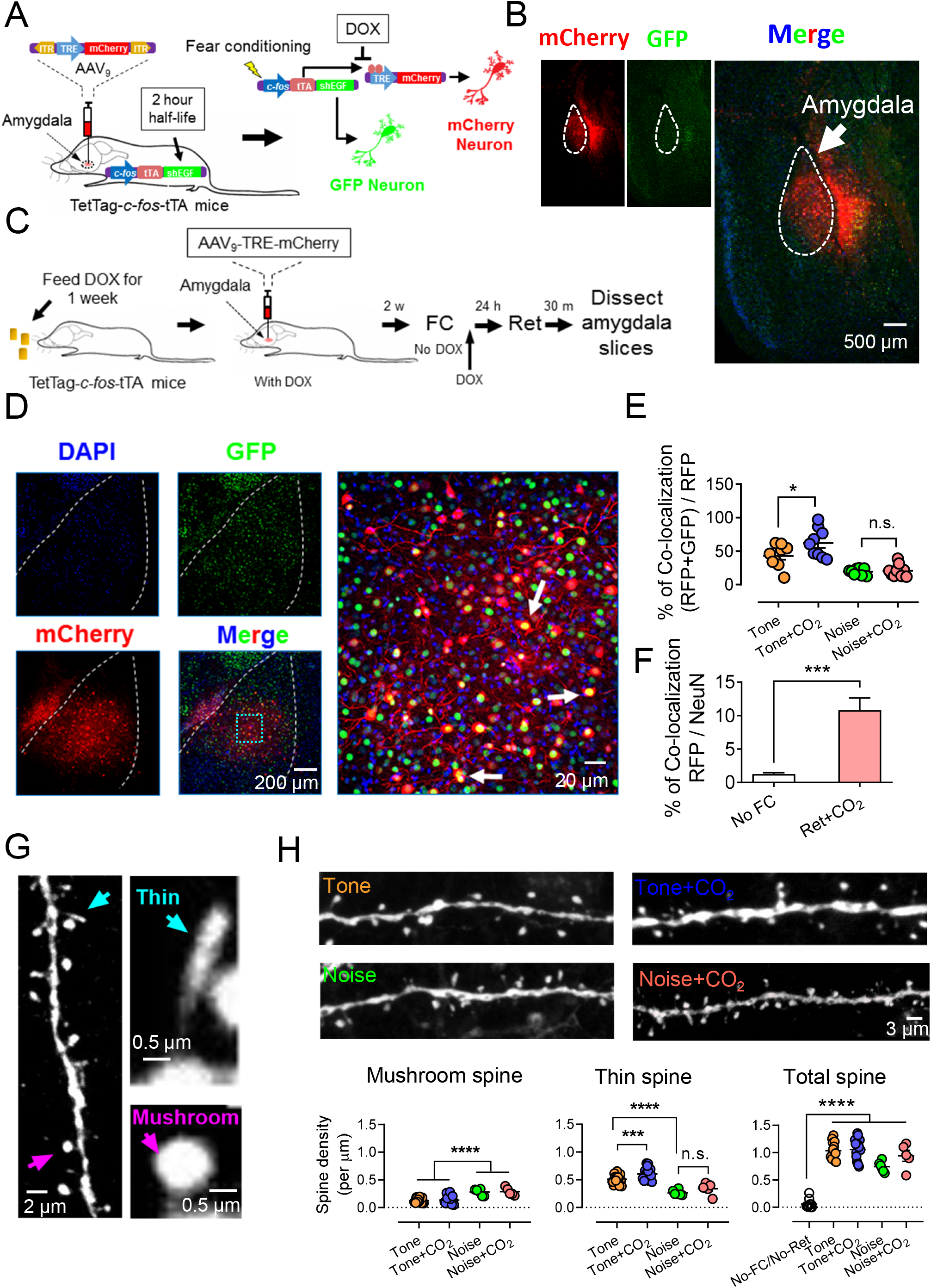
CO_2_ inhalation during a selective memory retrieval enhances the retrieval-related memory trace. **(A)** Schematic showing the c-Fos-tTA-GFP mouse system combined with an AAV_2/9_-mCherry to label a specific memory trace. The Fos promoter in transgenic mice was activated by activities, followed by a transient, 2-hour half-life, GFP expression in the cells. When the AAV_2/9_-mCherry virus was injected into the brain, the activation of c-FOS also induces the expression of mCherry. When the mice were fed with DOX, the mCherry expression was interrupted. **(B)** an example image showing the efficiency of the expression of GFP and mCherry in the amygdala. **(C)** The procedure of aversive conditioning, memory retrieval, and memory trace labeling using the system in A. Mice was fed with DOX for at least one week, followed by an injection of AAV_2/9_-mCherry in the amygdala. Two weeks later, DOX was removed and mice were subjected to aversive conditioning with a pure tone as the CS. DOX has added back again immediately after aversive conditioning. 24 hours later, the mice underwent retrieval with pure tone or noise. The brain slices were collected 30 mins after retrieval and then were stained for microscopy. **(D)** Left, representative images of the neurons labeled by mCherry, GFP, and DAPI; Right, the enlarged area from the “merge” image showing the overlapping expression of mCherry and GFP neurons. The overlapping neurons indicate their “consanguinity” in the same memory trace. **(E)** Summarized data are the percentage of the overlapping expression of mCherry and GFP neurons in different behavior groups. All mice underwent aversive conditioning with a tone as the CS. 24 hours later, the mice were separated into 4 groups. Tone group: retrieval CS by tone; Tone+CO_2_ group: retrieval CS by tone along with CO_2_ inhalation; Noise group: retrieval CS by noise; Noise +CO_2_ group: retrieval CS by noise along with CO_2_ inhalation. **(F)** Control experiment showing the expression of mCherry with or without DOX as well as with or without aversive conditioning. **(G)** Left, a representative image showing the spine morphology in the mCherry and GFP colocalized neurons. The mature spines were categorified as “mushroom” spines and the immature spines were categorified as “thin” spines; Right, an enlarged image showing the details of mushroom and thin spines. **(H)** Upper, representative images of the spine structures in different animal groups showed in E; Lower, summarized data of the spine densities of mushroom spines, thin spines, and total spines in the different groups. Data are mean ± SEM. n =8-12 mice for each group. ‘n.s.’ indicates not statistically significant. * indicates p<0.05, *** indicates p<0.001, **** indicates p<0.0001,by unpaired Student’s t-test and one-way ANOVA with Tukey’s post-hoc multiple comparisons.

We then examined the effects of CO_2_ on dendritic spine morphology after memory retrieval. Spine morphology has been widely indicated in the mechanism of synaptic plasticity (Woolfrey and Srivastava, 2016; Wright et al., 2020; Yang et al., 2009). Dendritic spines are the primary target of neurotransmission input in the central nervous system (Bourne and Harris, 2008), and their density and structure provide the basis for physiological changes in synaptic efficacy that underlie learning and memory (Bailey et al., 2015). Spine formation and plasticity are regulated by many conditions, including exterior stimulation and behavior (Gipson and Olive, 2017). We hypothesized that CO_2_ inhalation during retrieval alters both structure and plasticity of dendritic spines. The molecular mechanism by which CO_2_ regulates spine plasticity may explain how CO_2_ converts memory into the labile stage.

Using the TetTag-c-fos driven-GFP mouse model, we imaged spine structure and assessed spine density and morphology in overlapping neurons of the amygdala in each group (pure tone aversive conditioning, pure tone retrieval and pure tone aversive conditioning, white noise retrieval) **(Fig. 7A, C)**. Mature spines - most of which display “mushroom-like” morphology - have more stable postsynaptic structures enriched in AMPARs. In contrast, immature spines with a “thin-like” morphology, are unstable postsynaptic structures that have the transitional ability. Immature dendritic spines are thought to be responsible for synaptic plasticity, as they have the potential for strengthening (Berry and Nedivi, 2017). The categories of spines were identified based on the parameters in the previous studies **(Fig. 7G)** (see the details in the Material and Methods) (Kreple et al., 2014; Wright et al., 2020). The behavior procedure was described in **Fig. 7C**, the animals underwent aversive conditioning with a tone as a CS and followed by a retrieval on day 2 with tone or noise. We found increased spine numbers after aversive conditioning, indicating that aversive conditioning increases synaptic strength. There was no additional increase in the density of spines in all groups, suggesting that retrieval and CO_2_ inhalation did not change the synaptic strength **(Fig. 7H, lower-right)**.

We further analyzed spine subtypes as described in the experimental procedures. Thin spines are deemed to represent the immature structure of the synapses (Berry and Nedivi, 2017). We examined the ratio of the number of thin spines to the total number of spines. Interestingly, we found that there were more thin spines in the retrieval group, which might suggest a greater potential for synaptic plasticity in this group **(Fig. 7H, lower-middle)**. Importantly, when the retrieval group (tone) was paired with CO_2_, we found an additional increase of thin spines compared to the retrieval group alone **(Fig. 7H, lower-middle)**. This finding suggests that CO_2_ paired with retrieval might boost synaptic plasticity compared to memory retrieval alone. Consistently, the mushroom spine numbers decreased in the tone and CO_2_ paired retrieval groups, suggesting a higher turnover rate after memory retrieval **(Fig. 7H, lower-left)**. However, when the mice were trained with the pure tone and during the retrieval session were presented with white noise (a generated unrelated CS), we found that in the noise group with or without CO_2_ inhalation, the thin spine number did not increase compared to the pure tone/retrieval group **(Fig. 7H, lower-middle)**. This finding supports the specific effect of CO_2_ on the memory trace.

## DISCUSSION

A newly acquired aversive memory is labile and can be easily disrupted before it is transformed into a long-term stable state (Alberini, 2011). An existing memory, when reactivated, may become labile again during a short post-reactivation period known are reconsolidation window (Schiller et al., 2012). Previous studies using auditory aversive conditioning found that a retrieval event utilizing a single tone CS renders the memory labile during the reconsolidation window (Monfils et al., 2009). During this reconsolidation window, memory is sensitive to the updating processes that may either enhance or weaken the original memory (Bouton et al., 2020; Du et al., 2017; Fukushima et al., 2014). The reconsolidation window offers an opportunity to determine the mechanisms underlying the lability of an existing memory. We have recently found that when mice inhale 10% CO_2_ during retrieval, memory lability increases, and the original memory may be replaced through the update mechanisms (Du et al., 2017). Here, we show that the effects of CO_2_ on memory are specific to the reactivated conditioned cue (Debiec et al., 2010; Doyere et al., 2007). To address the specificity of the effects of CO_2_ on reactivated memories, we used a unique behavioral approach with distinct CSs and US, which allowed us the creation of distinct memories. An application of CO_2_ to the reactivated but not no-reactivated memory enhanced its lability.

We then asked about the cellular mechanisms through which CO_2_ enhances the lability of aversive memory. Memory retrieval-induced lability is glutamate receptor-dependent (Rose and Rankin, 2006; Milton, A.L., Everitt, B.J. 2013). We have shown that CO_2_ activates ASIC1a by decreasing pH in the brain during retrieval and the activation of ASIC1a increases postsynaptic intracellular calcium and increases AMPAR exchange (Du et al., 2017). Previous research studying the mechanism of retrieval of aversive memories have revealed the rapid and transient exchange from CI-AMPARs to CP-AMPARs in the lateral amygdala synapses (Clem and Huganir, 2010; Hong et al., 2013) after presenting the CS. In this study, we focused on the specificity of CO_2_ effects on the exchange of the AMPARs in the lateral amygdala’s synapses. We conditioned the mice with a pure tone CS and reactivated the memory with the pure tone with or without CO_2_ inhalation. Consistent with our earlier findings, we observed that CO_2_ increases AMPAR exchange when it is paired with retrieval. However, when mice were presented with an unrelated CS during the retrieval stage, CO_2_ did not alter the AMPAR exchange-suggesting the effects of CO_2_ on memory trace are memory specific. In addition, synaptic strength (ratio of AMPAR/NMDAR and amplitude of the mEPSCs) was not altered while applying CO_2_ during retrieval, with or without combining with the memory trace. The exchange of CI-AMPARs to CP-AMPARs indicates a synaptic plasticity change. Interestingly, CO_2_ did not increase the total number of spines compared to the retrieval alone group, which suggests the strength of the synapse in a memory trace does not change. In addition, thin spine density significantly increased when retrieval was combined with CO_2_ inhalation, suggesting that CO_2_ application changes plasticity. Using this model, we found when an unrelated CS was presented during the retrieval session, no additional increase of immature spine density occurred. This finding indicates no plasticity change in the memory trace. Thus, we can conclude from our findings that the effects of CO_2_ on memory trace are specific.

Although we have provided evidence that CO_2_ indeed acts with specificity on a memory trace, this study did not determine how CO_2_ directly regulates a memory. We cannot exclude the possibility that CO_2_ might trigger specific effects on memory through other targets. For instance, CO_2_ inhalation increases cerebral blood flow and arterial blood pressure and might affect brain functions, such as cognition. Although no direct evidence supports the possibility that increased cerebral blood flow and arterial blood pressure affect learning and memory, future studies will have the mechanisms underlying the specificity of CO_2_ on memory. Future studies will also have to examine whether CO_2_ has similar effects on synapses in other brain regions and on other learned behaviors (*e.g*. appetitive behaviors and drug addictive behaviors).

In conclusion, the effects of CO_2_ on the lability of aversive memory are found to be specific under certain conditions. Our research tests the novel hypothesis that protons are neurotransmitters that activate the postsynaptic proton receptors, ASICs, to manipulate memory updates. This non-invasive, drug-free methodology is innovative, efficacious, and safe for translation to clinical use. As a result, this research may lead to an effective complementary treatment for many mental health-related disorders for which efficient treatments are lacking. We hope this research will lead to new areas of inquiry into CO_2_-related mechanisms that underlie memory modification and lead to the development of novel therapies for anxiety disorders such as PTSD.

## ACKNOWLEDGMENTS

We thank Olivia Miller, Melissa Curtis, Nora Abdul-Aziz, Rida Naqvi, Caitlin Kilmurry, Becca Sturges, Jen Page, Chase Carr, Jordan Jones for their assistance. We thank Drs. Susumu Tonegawa for providing the TRE-mCherry plasmid. J.Du. is supported by the National Institutes of Mental Health (1R01MH113986), the University of Toledo start-up fund, and the University of Tennessee Health Science Center start-up fund.

## AUTHOR CONTRIBUTIONS

J.DU., J.D., E.K., C.K. and conceived the project. J.DU., E.K., C.K. and J.D. designed the experiments. E.K., C.K., J.E., S.B., M.H., B.L., performed the behavior experiments. K.S., performed the spine morphology experiments and data analysis. F.N., and J.DU. performed the patch-clamp experiments and data analysis. E.K., and J.DU., wrote the manuscript. All authors reviewed and edited the manuscript.

---

**Figure s1:**
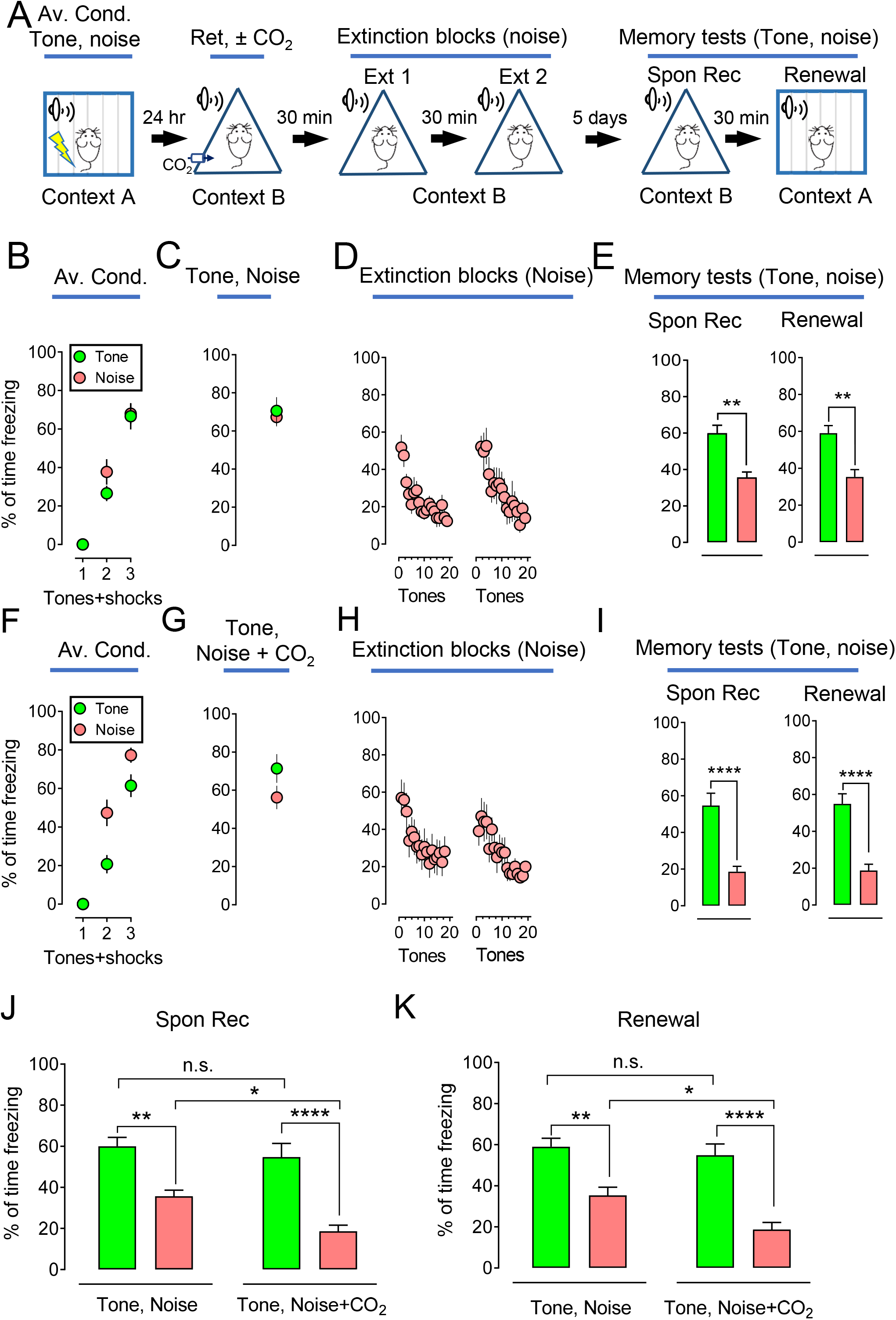
White noise was an appropriate successful CS to involve use in studying the effects of CO_2_ effects on aversive conditioning and retrieval. **(A)** Representative schematic of protocol for the aversive conditioning (pure tone and white noise), memory retrieval (pure tone and white noise), extinction (white noise), and memory test (spontaneous recovery and renewal). **(B-E)** Data are presented by the percentage of freezing time during the CSs (tone and noise) in aversive conditioning **(B)**, retrievals (tone and noise) **(C)**, two sections of extinction with white noise **(D)**, memory test of spontaneous recovery (Spon Rec) and renewal with tone and noise **(E)**. **(F-I)** Data are presented by the percentage of freezing time during the CSs (tone and noise) in aversive conditioning **(F)**, retrievals (pure tone and white noise plus CO_2_ inhalation) **(G)**, two sections of extinction with white noise **(H)**, memory test of spontaneous recovery (Spon Rec) and renewal with tone and noise **(I)**. **(J-K),** comparison data based on spontaneous recovery and renewal respectively from panels **E** and **I**. Data are mean ± SEM. n = 12 mice in each group. ‘n.s.’ indicates not statistically significant. * indicates p<0.05, ** indicates p<0.01, *** indicates p<0.001, *** indicates p<0.0001, by unpaired Student’s t-test and one-way ANOVA with Tukey’s post-hoc multiple comparisons.

**Figure s2:**
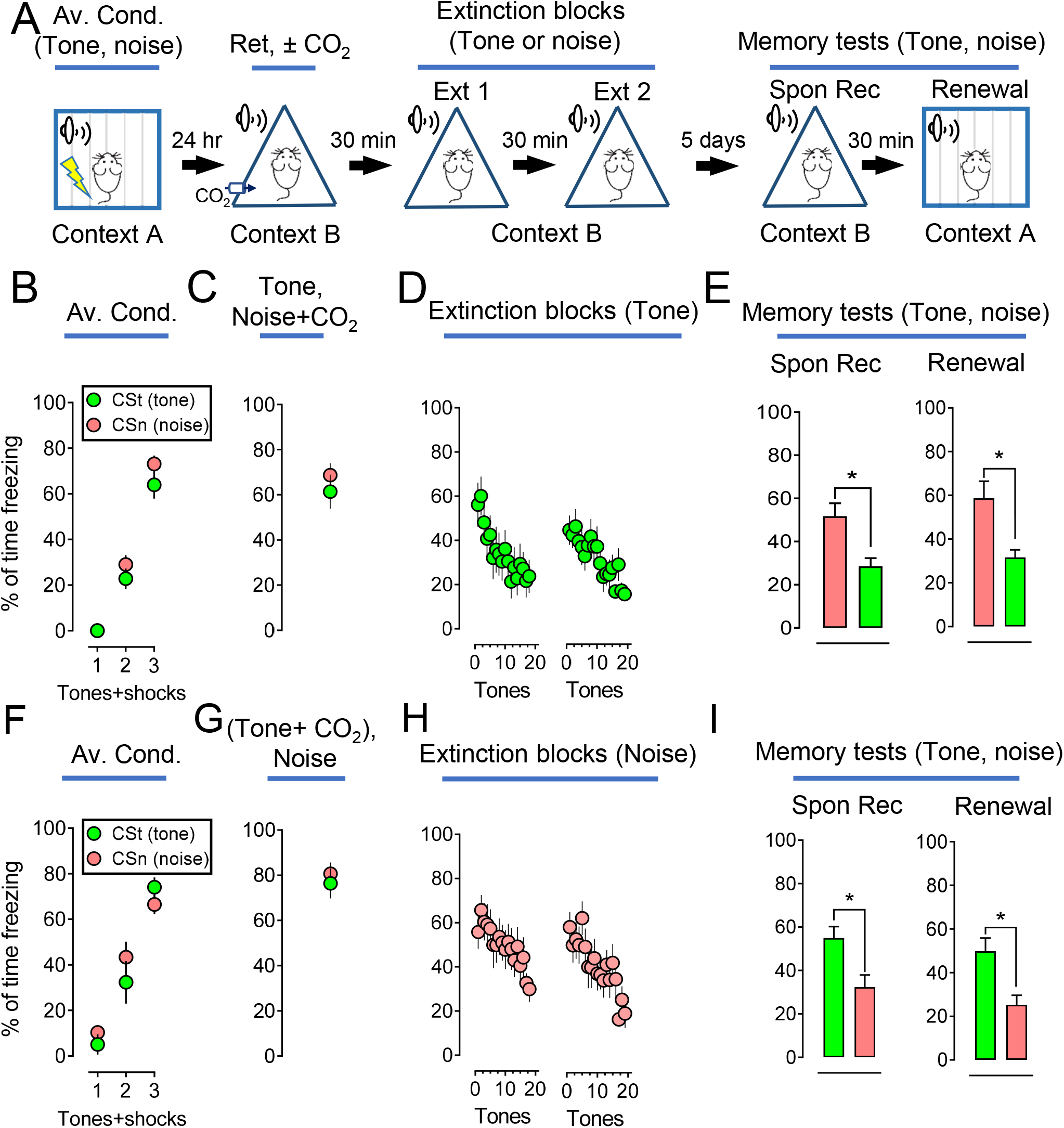
CO_2_ inhalation does not boost extinction when pairs with an unrelated CS during memory retrieval. **(A)** Representative schematic of protocol for the aversive conditioning (pure tone and white noise), memory retrieval (pure tone and white noise), extinction (white noise), and memory test (spontaneous recovery and renewal). **(B-E)** Data are presented by the percentage of freezing time during the CSs (tone and noise) in aversive conditioning **(B)**, retrievals (pure tone and white noise plus CO_2_ inhalation) **(C)**, two sections of extinction with pure tone **(D)**, memory test of spontaneous recovery (Spon Rec) and renewal with tone and noise, Blue arrow and % indicated the difference (decreases) between the tone and noise groups **(E)**. **(F-I)** Data are presented by the percentage of freezing time during the CSs (tone and noise) in aversive conditioning **(F)**, retrievals (pure tone plus CO_2_ inhalation and white noise) **(G)**, two sections of extinction with white noise **(H)**, memory test of spontaneous recovery (Spon Rec) and renewal with tone and noise, Blue arrow and % indicated the difference (decreases) between the tone and noise groups. **(I)**. Data are mean ± SEM. n = 12 mice in each group. * indicates p<0.05 by unpaired Student’s t-test.

**Figure s3:**
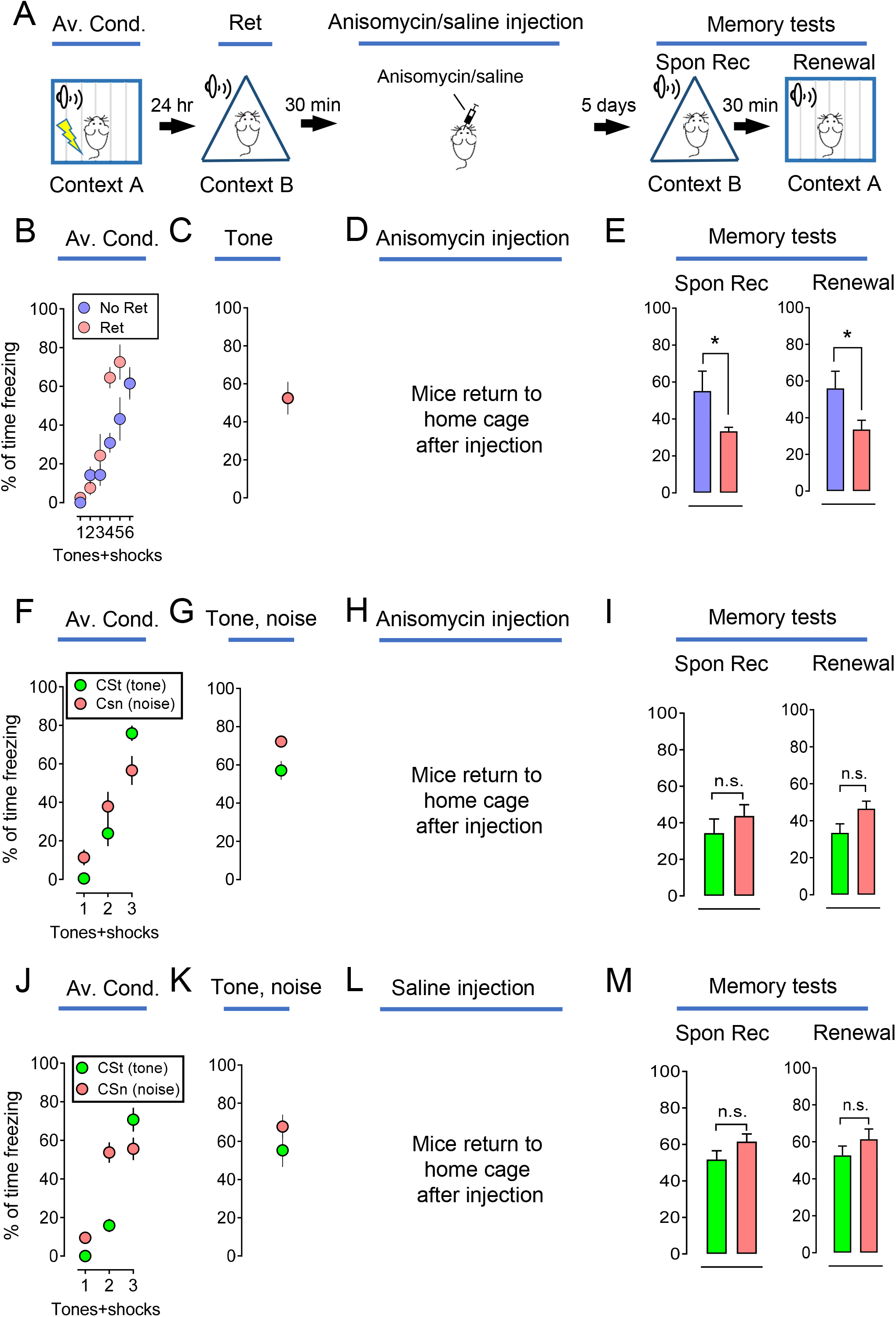
Anisomycin disrupts the memory within the reconsolidation window. **(A)** Representative schematic of protocol for the aversive conditioning (pure tone), with or without memory retrieval (pure tone), anisomycin infusion, and memory test (spontaneous recovery and renewal). **(B-E)** Data are presented by the percentage of freezing time during the tone presentation in aversive conditioning **(B)**, with or without retrieval (tone) **(C)**, anisomycin infusion in the amygdala **(D)**, spontaneous recovery (Spon Rec), and renewal test with tones **(E)**. **(F-I)** Data are presented by the percentage of freezing time during the CSs (tone and noise) in aversive conditioning **(F)**, retrievals (pure tone and white noise) **(G)**, anisomycin injection in the amygdala **(H)**, memory test of spontaneous recovery and renewal with tone and noise **(I)**. **(J-M)** Data are presented by the percentage of freezing time during the CSs (tone and noise) in aversive conditioning **(J)**, retrievals (pure tone plus CO_2_ inhalation and white noise) **(K)**, saline injection in the amygdala **(L)**, memory test of spontaneous recovery and renewal with tone and noise, **(M)**. Data are mean ± SEM. n = 11-12 mice in each group. ‘n.s.’ indicates not statistically significant. * indicates p<0.05 by unpaired Student’s t-test.

**Figure s4:**
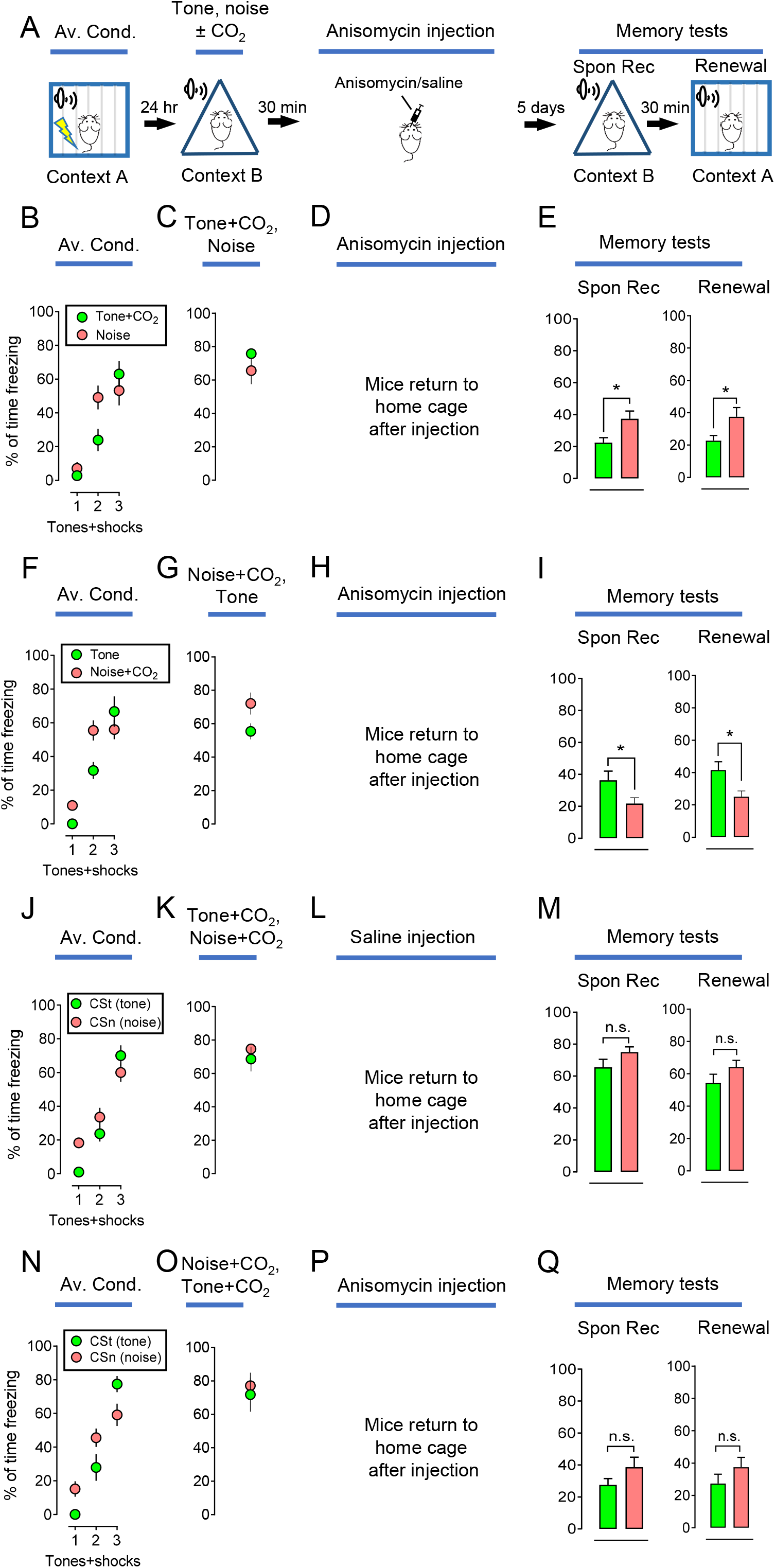
CO_2_ enhances memory retrieval and potentiates the effects of anisomycin on the reconsolidation. **(A)** Representative schematic of protocol for the aversive conditioning (pure tone and white noise), memory retrieval (pure tone and white noise) with or without CO_2_ inhalation, anisomycin infusion, and memory test (spontaneous recovery and renewal). **(B-E)** Data are presented by the percentage of freezing time during the CSs (tone and noise) presentation in aversive conditioning **(B)**, retrieval (tone plus CO_2_ and noise) **(C)**, anisomycin infusion in the amygdala **(D)**, spontaneous recovery (Spon Rec) and renewal test with tone and noise **(E)**. **(F-I)** Data are presented by the percentage of freezing time during the CSs (tone and noise) in aversive conditioning **(F)**, retrieval (noise plus CO_2_ and tone) **(G)**, anisomycin infusion in the amygdala **(H)**, memory test of spontaneous recovery (Spon Rec) and renewal with tone and noise **(I)**. **(J-M)** Data are presented by the percentage of freezing time during the CSs (tone and noise) in aversive conditioning **(J)**, retrievals (tone and noise) plus CO_2_ **(K)**, saline infusion in the amygdala **(L)**, memory test of spontaneous recovery (Spon Rec) and renewal with tone and noise **(M)**. **(N-Q)** Data are presented by the percentage of freezing time during the CSs (tone and noise) in aversive conditioning **(N)**, retrievals (tone and noise) plus CO_2_ **(O)**, anisomycin infusion in the amygdala **(P)**, memory test of spontaneous recovery (Spon Rec) and renewal with tone and noise **(Q)**. Data are mean ± SEM. n = 12 mice in each group. ‘n.s.’ indicates not statistically significant. * indicate p<0.05 by unpaired Student’s t-test.

**Figure s5:**
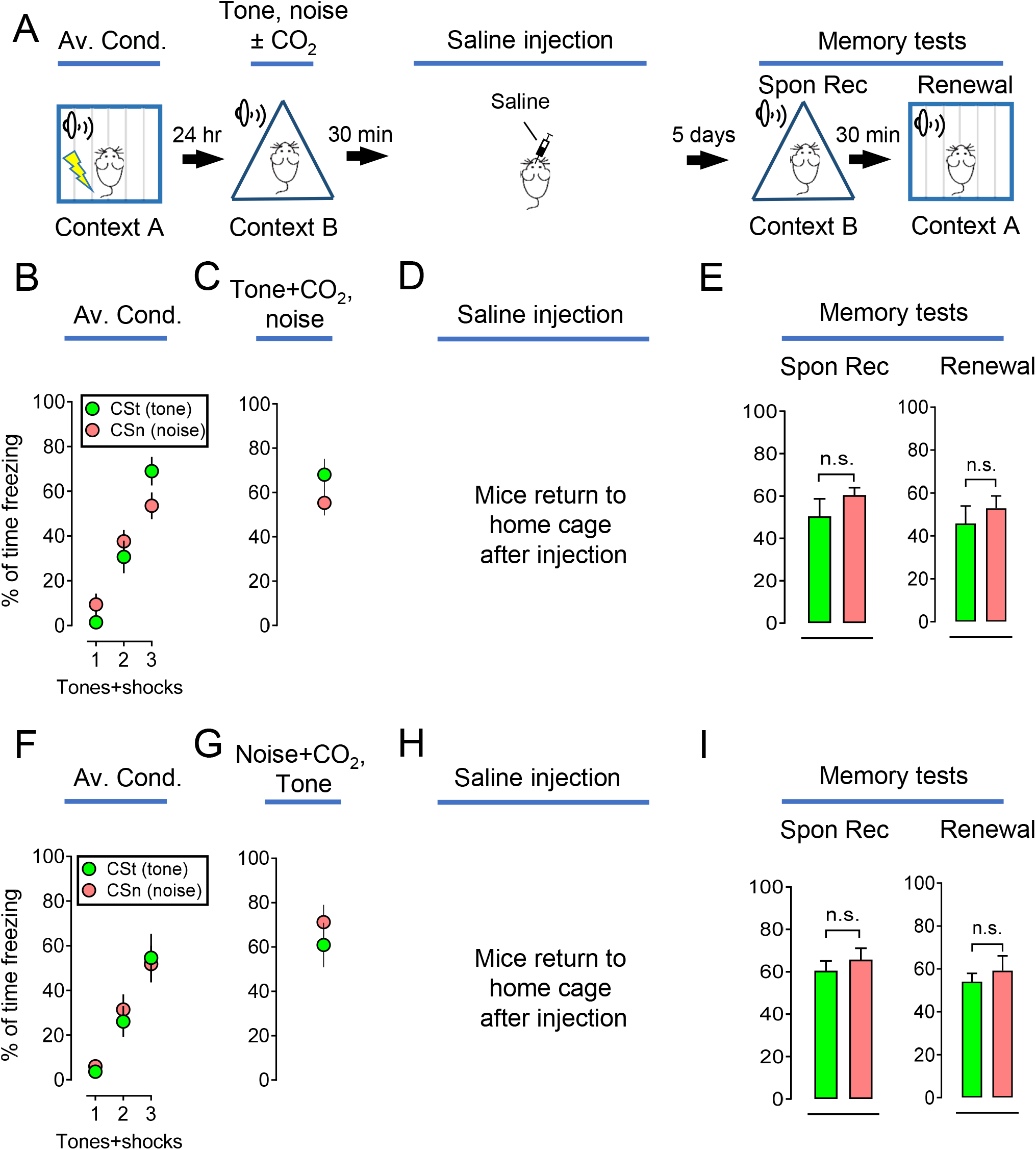
CO_2_ does not affect memory reconsolidation without anisomycin. **(A)** Representative schematic of protocol for the aversive conditioning (pure tone and white noise), memory retrieval (pure tone and white noise) with or without CO_2_ inhalation, saline infusion, and memory test (spontaneous recovery and renewal). **(B-E)** Data are presented by the percentage of freezing time during the CSs (tone and noise) presentation in aversive conditioning **(B)**, retrieval (tone plus CO_2_ and noise) **(C)**, saline infusion in the amygdala **(D)**, spontaneous recovery (Spon Rec) and renewal test with tone and noise **(E)**. **(F-I)** Data are presented by the percentage of freezing time during the CSs (tone and noise) in aversive conditioning **(F)**, retrieval (noise plus CO_2_ and tone) **(G)**, saline infusion in the amygdala **(H)**, memory test of spontaneous recovery (Spon Rec) and renewal with tone and noise **(I)**. Data are mean ± SEM. n = 12 mice in each group. ‘n.s.’ indicates not statistically significant by unpaired Student’s t-test.

